# Comprehensive Analysis of Ubiquitously Expressed Genes in Human, From a Data-Driven Perspective

**DOI:** 10.1101/2021.02.09.430465

**Authors:** Jianlei Gu, Jiawei Dai, Hui Lu, Hongyu Zhao

**Author notes:** Correspondence: Hui Lu, Hongyu Zhao.

## Abstract

Comprehensive characterization of spatial and temporal gene expression patterns in humans is critical for uncovering regulatory codes of the human genome and understanding molecular mechanisms of human diseases. The ubiquitously expressed genes (UEGs) refer to those genes expressed across a majority, if not all, phenotypic and physiological conditions of an organism. It is known that many human genes are broadly expressed across tissues. However, most previous UEG studies have only focused on providing a list of UEGs without capturing their global expression patterns, thus limiting the potential use of UEG information. In this article, we propose a novel data-driven framework to leverage the extensive collection of ∼40,000 human transcriptomes to derive a list of UEGs and their corresponding global expression patterns, which offers a valuable resource to further validate and characterize human UEGs. Our results suggest that about half (12,234; 49.01%) of the human genes are expressed in at least 80% of human transcriptomes and the median size of the human transcriptome is 16,342 (65.44%). This suggests that the average difference in gene content between human transcriptomes is only 16.43%. Through gene clustering, we identified a set of UEGs, named LoVarUEGs, that have stable expression across human transcriptomes and can be used as internal reference genes for expression measurement. To further demonstrate the usefulness of this resource, we evaluated the uniqueness of repression for 16 previously predicted disallowed genes in islets beta cells and found that seven of these genes showed relatively more varied expression patterns, suggesting that the repression of these genes may not be unique to islets beta cells. We have made our resource publicly available at https://github.com/macroant/HumanUEGs.

## Introduction

In multicellular organisms, different tissues or cells contain mostly the same genome. However, each tissue or cell type only expresses a subset of its genes and has its own unique transcriptome. The variations among transcriptomes underlie the wide range of phenotypic and physiologic differences across tissues or cells [1]. It is generally believed that the genes within a transcriptome could be broadly divided into two groups: the ubiquitously expressed genes (UEGs), traditionally called housekeeping genes (HKs) [2], and the specifically expressed genes (SEGs) [3]. UEGs are expressed in almost all living cells of an organism and play an essential role in maintaining cellular processes and cell survival. On the other hand, SEGs are strictly expressed in a limited number of tissue or cell types and usually have specific biological functions. They are generally believed to be more likely associated with human diseases and/or druggable targets [4]. The more recent view of UEGs or HKs has emphasized that these genes should be insensitive to cell type heterogeneity and have stable expression across tissues [5, 6]. In this study, we use the term UEGs rather than HKs to describe those widely expressed genes with some having variations across conditions, and systematically characterize the global expression patterns of UEGs in the human genome.

Much work has been conducted to characterize the UEGs in the human genome. However, the reproducibility of the UEG lists from early studies was low due to the limitations of microarray techniques [6]. As far as we know, it was not until 2008 that Jiang et.al [7] first reported that there might be a large number of human genes (about 40%) broadly expressed across tissues through the analysis of an EST (Expressed Sequence Tag) data collection. With the development of the RNA-seq technology, this observation was substantiated by RNA-seq studies with approximately 8,000 to 10,000 genes broadly expressed across tissues [8, 9]. However, there are several limitations in the published UEG studies. First, there are over 200 tissue/cell types in the human body and there can be substantial variations in transcriptomes across biological conditions and individuals [10, 11]. The published UEG studies are often limited in the number of tissue and cell types covered. Second, published UEG studies often use a single tissue-specificity measure of expression to identify UEGs and do not fully capture gene expression patterns, thus limiting the potential use of UEG information. Although some UEG studies have considered expression variability, it has only been used as a hard filtering criterion [5]. For bulk RNA-seq data, the observed expression level for each gene is the aggregated expression value of a large number (maybe heterogeneous) of cells. Thus, traditional bulk RNA-seq data offer a bird’s-eye view of the expression patterns at the cell population level.

Inspired by the concept of pan-genome and core-genome in bacterial research [12, 13], we hereby propose a novel analysis framework to systematically characterize human UEGs, which represents the core component of human transcriptomes. Through simultaneous consideration of a large collection of diverse transcriptomes, our framework bypasses the subjective tissue/cell type stratification process to directly assess the global expression specificity and the expression pattern for each gene (Fig 1). By analyzing ∼40,000 divergent human transcriptomes, we observed that 12,234 human genes (49.01%) are ubiquitously expressed in at least 80% of human transcriptomes. Coupled with global expression patterns of these genes, we identified a set of UEGs, named LoVarUEGs, that have stable expression across biological conditions and can be used as internal reference genes for expression measurement. We observed similar results in another RNA-seq data repository, supporting the generalizability of our findings. To demonstrate the usefulness of our UEG resource, we evaluated the global expression patterns of 16 previously predicted disallowed genes in pancreatic islets beta cells and found that seven of these putative disallowed genes have a more varied expression pattern than classical disallowed genes, suggesting that the repressions of these genes may not be unique to islet beta cells, at least in term of expression level. In summary, our study provides a useful framework and resource for further functional genomics studies of human genes.

**Fig 1.**
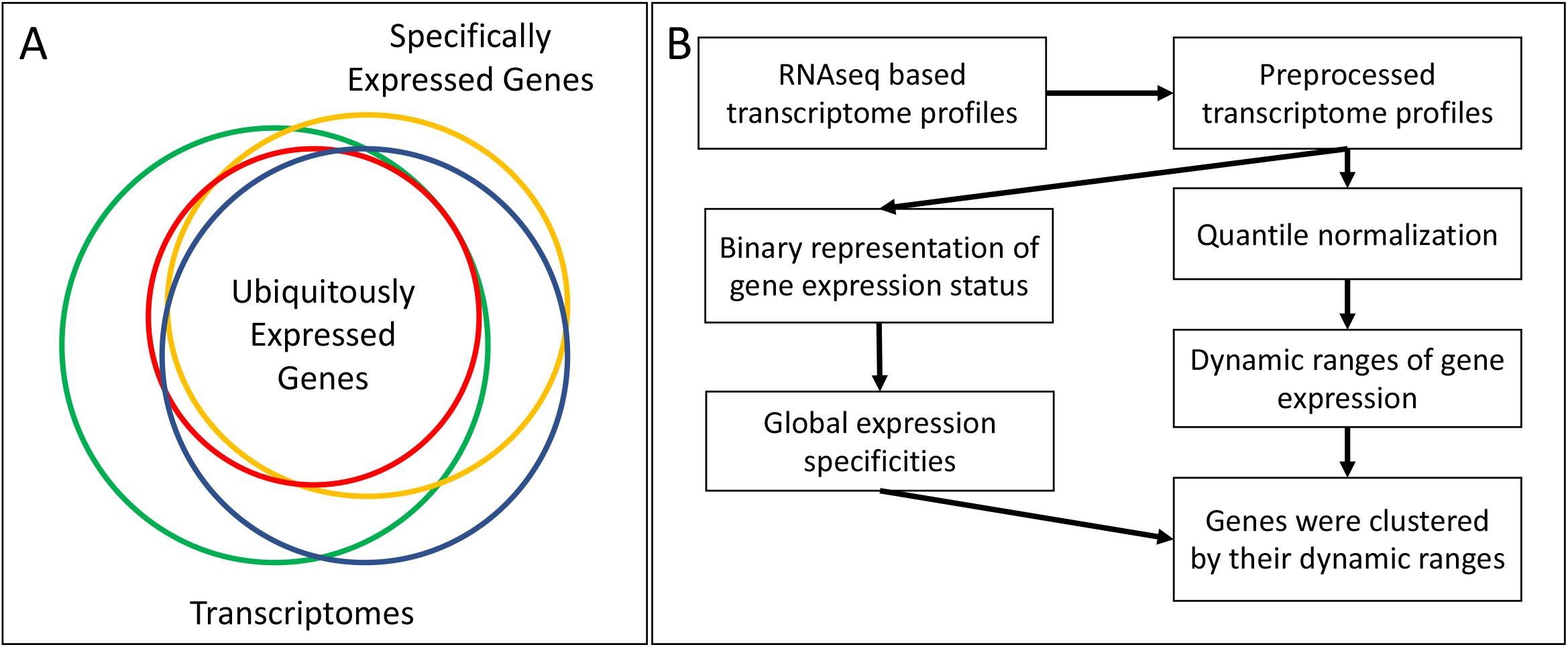
The flow diagram for systematic characterization of ubiquitously expressed genes in the human genome. (A) Definition of global expression specificity, different colored circles represent the transcriptomes derived from different tissues or cell types. The overlapping area represents the core component of human transcriptomes, the ubiquitously expressed genes (UEGs). (B) After preprocessing, we performed a sample-wise quantile normalization that allowed us to obtain a robust global distribution of expression level for each gene. Then, we clustered genes by their dynamic range of global distribution. Finally, the global expression specificity metrics were mapped to the genes and gene clusters.

## Results

### Summary of Analyzed Transcriptomes

In this study, we primarily use the recount2 repository [14, 15], which comprises ∼50,000 RNA-seq based human transcriptome profiles. After pre-processing (described in the Method section), 39,863 (80.3%) transcriptome profiles were retained for further analysis. We annotated the sample and tissue types of these transcriptomes with an automated semantic annotation database [16]. These transcriptomes covered more than 30 organ systems, with musculoskeletal system (10.09%) being the most common tissue, followed by hemolymphoid system (8.86%), nervous system (7.61%), and digestive system (2.74%) (Supp Table 2). To improve sample coverage across more conditions, we also included the transcriptomes from in vitro cells (including cell lines, primary cells, in vitro differentiated cells, stem cells, and iPS cells), with about 56.7% of the total samples (Supp Table 1). For reference, we further manually annotated 6,501 (16.31% of total) transcriptomes that represent 101 major tissue types (Supp Table 6). To check the relatedness among these transcriptomes, we used online PCA to visualize the first two principal components of all 39,863 transcriptome profiles (Fig 2). We can see that these divergent transcriptomes collected from various experiments are reasonably clustered, and those unclassified transcriptomes exhibited a broad transcriptomic heterogeneity. We found that 17,503 (43.91%) transcriptomes showed relatively high relatedness (Supp Fig 2), suggesting that these transcriptomes may be overrepresented in the recount2 dataset. We then conducted a sensitivity analysis to evaluate the impact of these overrepresented samples (described in the supplementary section) and observed that these overrepresented samples have limited effect on our overall results and conclusions. To evaluate the generalizability of our results, we applied our analysis framework to a more recent transcriptome dataset, Dee2[17], which is a public repository of uniformly processed RNA-seq profiles. The differences between Dee2 and Recount2 dataset are that 1) they used different pipelines to generate transcriptome profiles; 2) they only shared about 15% of the samples and had different relatedness patterns (Supp Fig 1 and Supp Fig 2).

**Fig 2.**
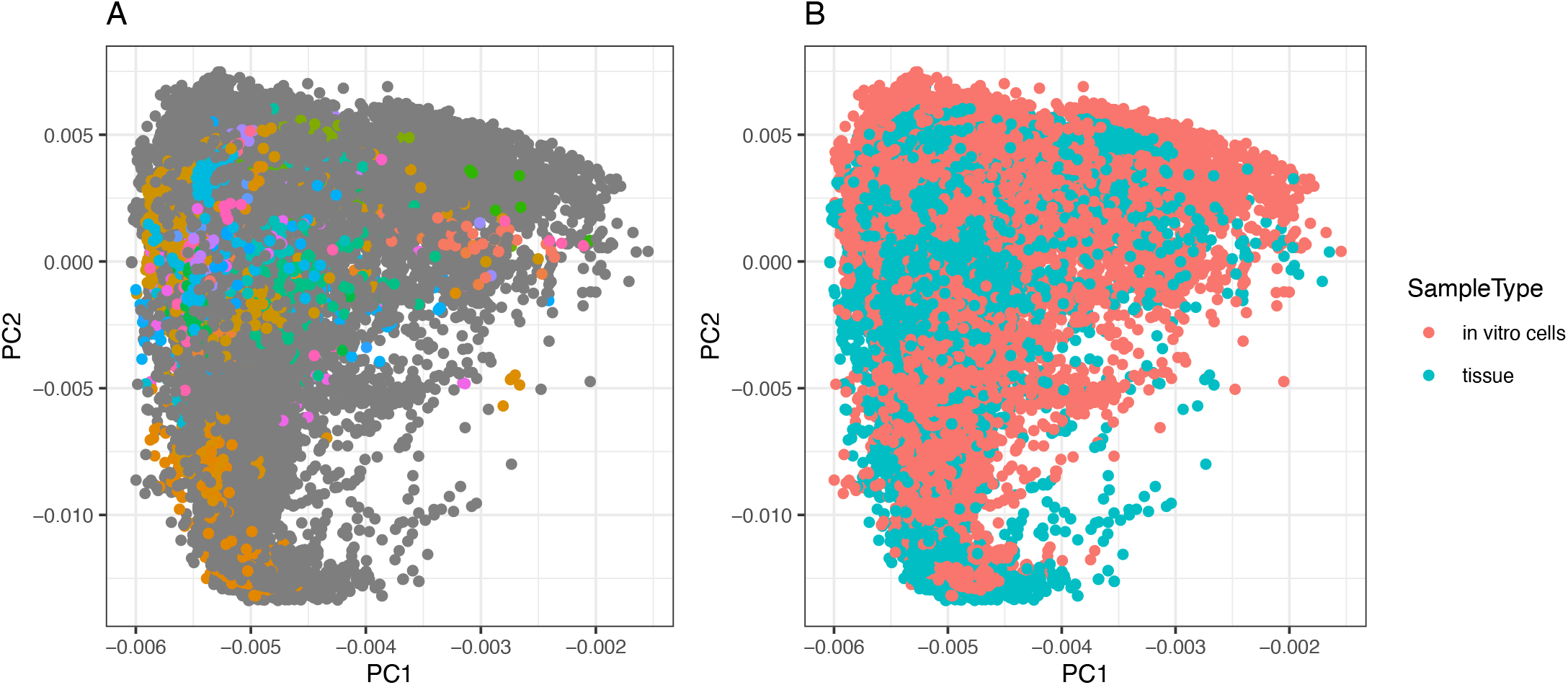
The phenotypic compositions and relatedness among analyzed transcriptomes. The onlinePCA was performed to the quantile normalized expression matrix to visualize the phenotypic compositions and relatedness among transcriptomes. Each dot represents one transcriptome projected on the principal plane formed by the first and second principal axes. (A). The colored dots represent the 6,501 (16.31%) manually curated reference transcriptomes belonging to 101 tissue groups. Dark dots represent those unclassified transcriptomes that also exhibited a wide spectrum of heterogeneity. (B). The blue dots are the transcriptomes from tissue samples, and the red dots are the transcriptomes from in vitro cells.

### Global Expression Specificity and the Size of Human Transcriptome

We propose to use the proportion of samples that a gene was expressed across all the transcriptomes to quantify its expression specificity. We refer to this proportion as global expression specificity ϕ, where ϕ=1 denotes a ubiquitously expressed gene and ϕ close to 0 denotes a highly specifically expressed gene. As shown in Fig 3C, the distribution of ϕ has a clear bimodal distribution, i.e., most genes are either ubiquitously or specifically expressed, which is consistent with previous observations [18]. In order to determine the optimal detection threshold, we made a comparison between four commonly used detection thresholds and found that TPM 0.1, which was used in the GTEx project, is a robust and sensitive expression detection threshold for those lowly-expressed genes (Supp Table 13 and Supp Fig 25). Applying this threshold, 12,267 (49.14%) genes had their ϕ >= 0.8 and 7,439 (29.80%) genes had ϕ <= 0.4 (Table 1). To compare ϕ with traditional tissue-based expression specificity using manually curated samples with tissue information, we calculated tissue-based specificity and compared these two metrics across genes. As shown in Supp Fig 3, these two metrics are highly correlated with a Pearson coefficient of 0.960. Only 2,279 genes (9.1%) had a difference >=0.2 (20% of total range) between these two specificity metrics, where genes with relatively high or low global expression specificity have higher agreement (Supp Fig 4). In addition, 80% of the human transcriptomes had between 11,166 (44.71%) and 19,033 (76.21%) expressed genes, and the median number of expressed genes is 16,342 (65.44%) (Fig 3B). This number is close to what was reported in a smaller scale study [8].

**Fig 3.**
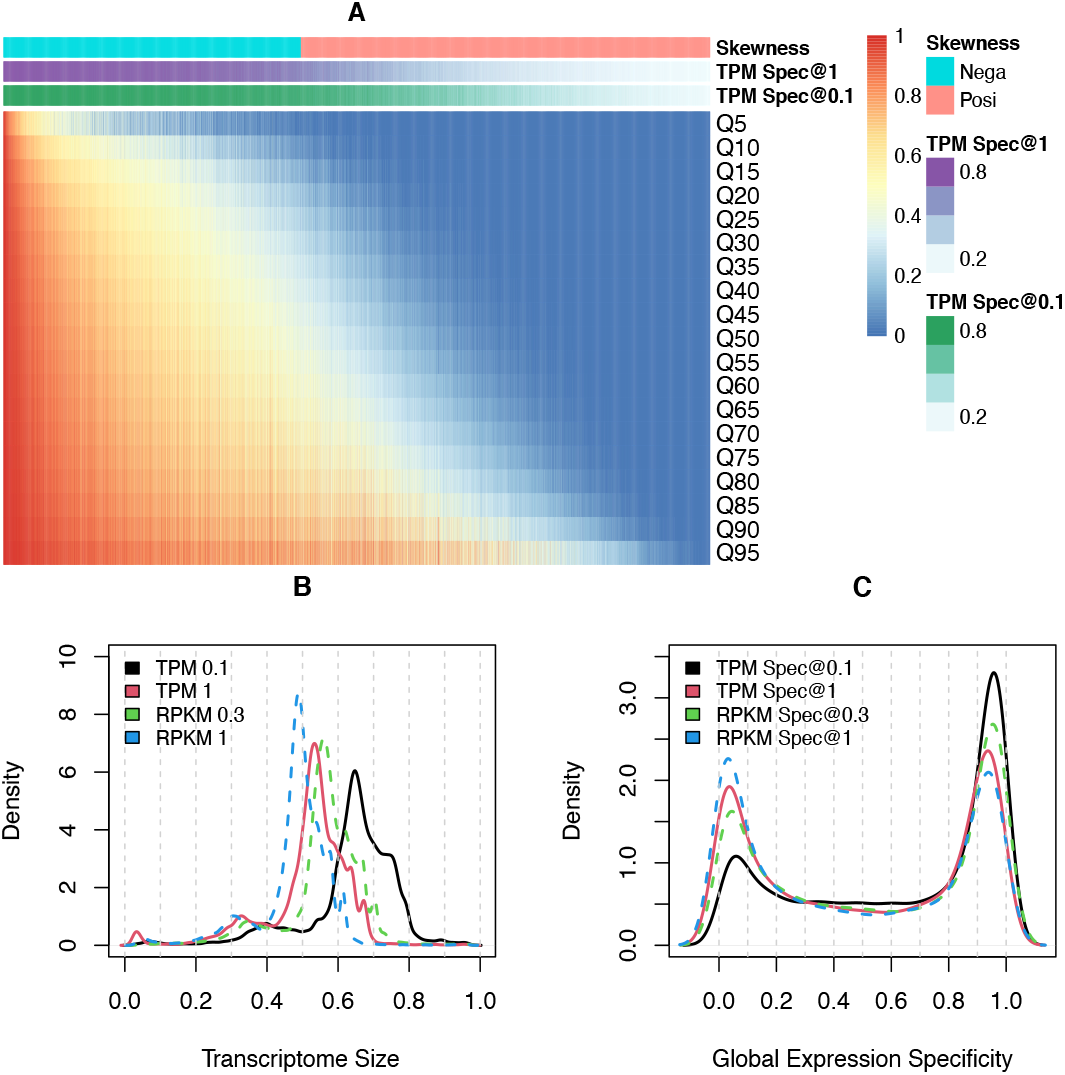
The global expression specificity and global dynamic ranges of expression values. (A) Quantile normalized transcriptome profiles summarized into dynamic ranges (the lowest 5% (Q5) to highest 95% (Q95) relative expression level), and used to generate a heatmap showing the global expression pattern for each gene. (B) The density plot shows the distribution of human transcriptome size. Apply the threshold of TPM 0.1, The median size of the human transcriptome is 16,342 (65.44% of human genes). (C) The density plot shows the global expression specificity obtained by different detection thresholds (1 TPM, 0.1 TPM, 1 RPKM, and 0.3 RPKM). It clearly shows that a large fraction of human genes are either ubiquitously expressed or specifically expressed. TPM Spec@1 and <u>Spec@0.1</u>donated the global expression specificity determined by TPM 1.0 and 0.1 threshold.

**Fig 4.**
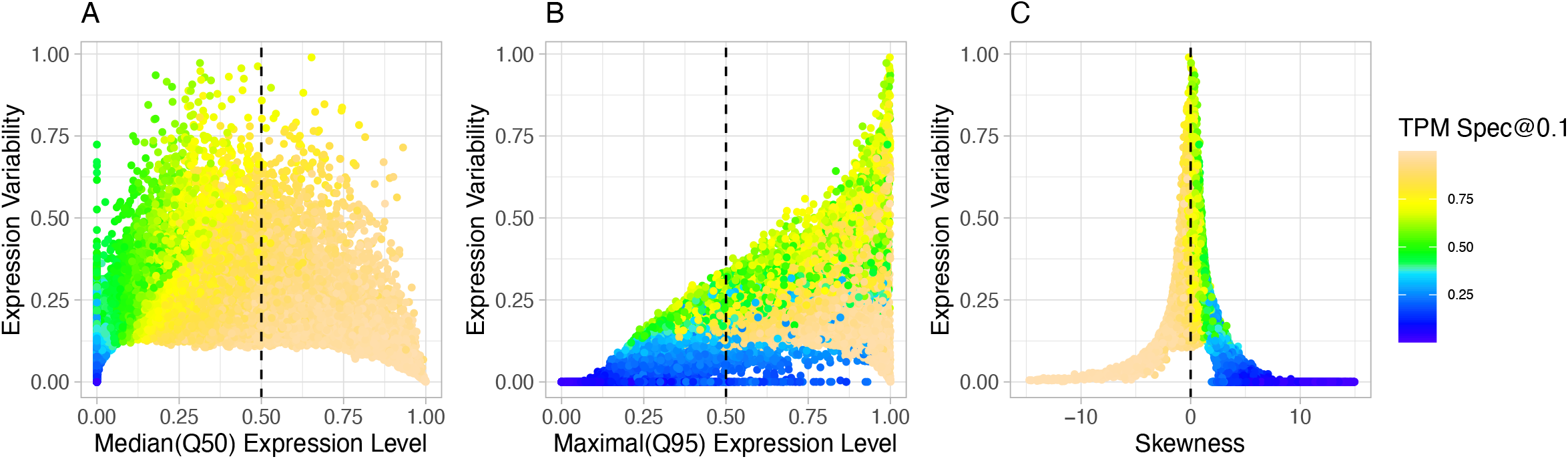
The relationships between expression specificity, expression level and expression variability. (A) and (B) show that UEGs tend to have higher median (Q50) and maximal (Q95) expression level. (C) The Scatter plot shows the relationship between distribution skewness, expression specificity, and expression variability. Global expression specificity of genes was highly correlated with their distribution skewness. UEGs tend to have negative distribution skewness, and SEGs tend to have positive distribution skewness. Most UEGs have relatively lower expression variability (IQR). TPM <u>Spec@0.1</u>donated the global expression specificity determined by 0.1 threshold.

### Global Expression Patterns of Human Genes

Although it is useful to classify genes into UEG and SEG groups by their expression specificity, such classification cannot capture global expression patterns. Through joint analysis of diverse transcriptomes, we can characterize the dynamic ranges of relative expression values for each gene. To reduce batch effects in defining global expression patterns, we used a sample-wise quantile transformation [19] to TPM (or RPKM) normalized transcriptome profiles. After transformation, expression values were replaced by their percentile ranks for each profile. Fig 3 (A) displays the dynamic ranges of gene expressions (the lowest 5% to the highest 95% relative expression values for each gene). With an empirical threshold of Q10 >= 0.1 (Supp Fig 23), the expression levels of 9,692 (38.83%) genes were above the 10% percentile in at least 90% of all transcriptomes. With a more relaxed threshold (Q20 >= 0.1), this number increased to 12,005 (48.09%), i.e. these genes’ expression levels were about the 10% percentile in at least 80% of the samples (Table 1). These observations are close to the inference of UEGs through ϕ. Supplementary Table 7 lists all the genes with their global expression specificity and dynamic ranges.

We then examined the relationships between global expression specificity and the distribution attributes of relative expression values, including mean, median, IQR (variability), and skewness. As expected, the genes with larger ϕ tended to have higher median expression levels (Fig 4), which is consistent with previous observations [18, 20, 21]. One interesting finding is that the global expression specificity is strongly associated with distribution skewness of relative expression values as observed in previous tissue specificity of gene expression (Spearman coefficient is −0.97) [18, 22]. The UEGs were enriched with genes showing negative-skewness, whereas the SEGs were enriched with genes having positive-skewness (Fig 3 A, Fig 4 C, and Table 1). About 90% of previously reported UEGs had a negatively-skewed distribution, and ∼80% of previously reported SEGs had a positively-skewed distribution.

### Global Expression Specificity Categories and Functional Implications

To group genes according to their global expression patterns, we performed clustering on the dynamic range matrix through percentile clustering [23], that is, to cluster genes according to their summarized distribution shapes of expression values. After clustering, the genes with similar expression levels, expression variability, and expression specificity were grouped into the same cluster (Fig 5). Figure 5A shows the PCA plot of the dynamic range matrix with 96 gene clusters inferred by the affinity propagation clustering method [24]. Fig 5B shows the dynamic range for some gene clusters, e.g., cluster #81 with most ubiquitously and highly expressed genes; cluster #10 with ubiquitously but lowly expressed genes; cluster #60 with the most varied expression pattern and cluster #70 with the most restricted expression pattern. We then mapped the global expression specificity to these gene-clusters, and with such information, we can broadly classify these gene clusters into five specificity categories:

1) UEG category (UEGs@1) detected by the threshold of TPM 1.0 (with median ϕ of clusters >= 0.8). This category included 9,687 (38.81%) genes in 40 clusters that had a ubiquitous expression pattern. The genes in this category are more likely involved in essential cellular processes, such as transcription (11.95%), apoptotic process (3.45%), oxidation-reduction process (3.28%), protein transport (3.16%), and cell division (2.95%).

2) UEG category (UEGs@0.1) only inferred by a more sensitive detection threshold of TPM 0.1 (with median ϕ of clusters >= 0.8). There are 14 clusters involving 2,547 (10.20%) genes in this category. Some clusters in this category showed low expression levels, e.g., cluster #77 and cluster #4, and they might be easily overlooked by stringent detection thresholds (Supp Table 13 and Supp Fig 25) or experiments with insufficient detection sensitivity. A total of 352 (78.40%) genes of these two gene clusters were above the 10% percentile in at least 80% of all transcriptomes studied, whereas only 4.45%-6.46% of them were classified as UEGs in previous UEG studies [8, 9]. On the other hand, some clusters in this category showed relatively higher expression variability, e.g., clusters #31, #65 and #55. Although these genes were widely detectable, their percentile ranks within each transcriptome varied significantly across biological conditions. This means that these gene clusters with higher expression variability are more likely to have a leaky expression [5, 25] or more sensitive to cell type heterogeneity.

**Fig 5.**
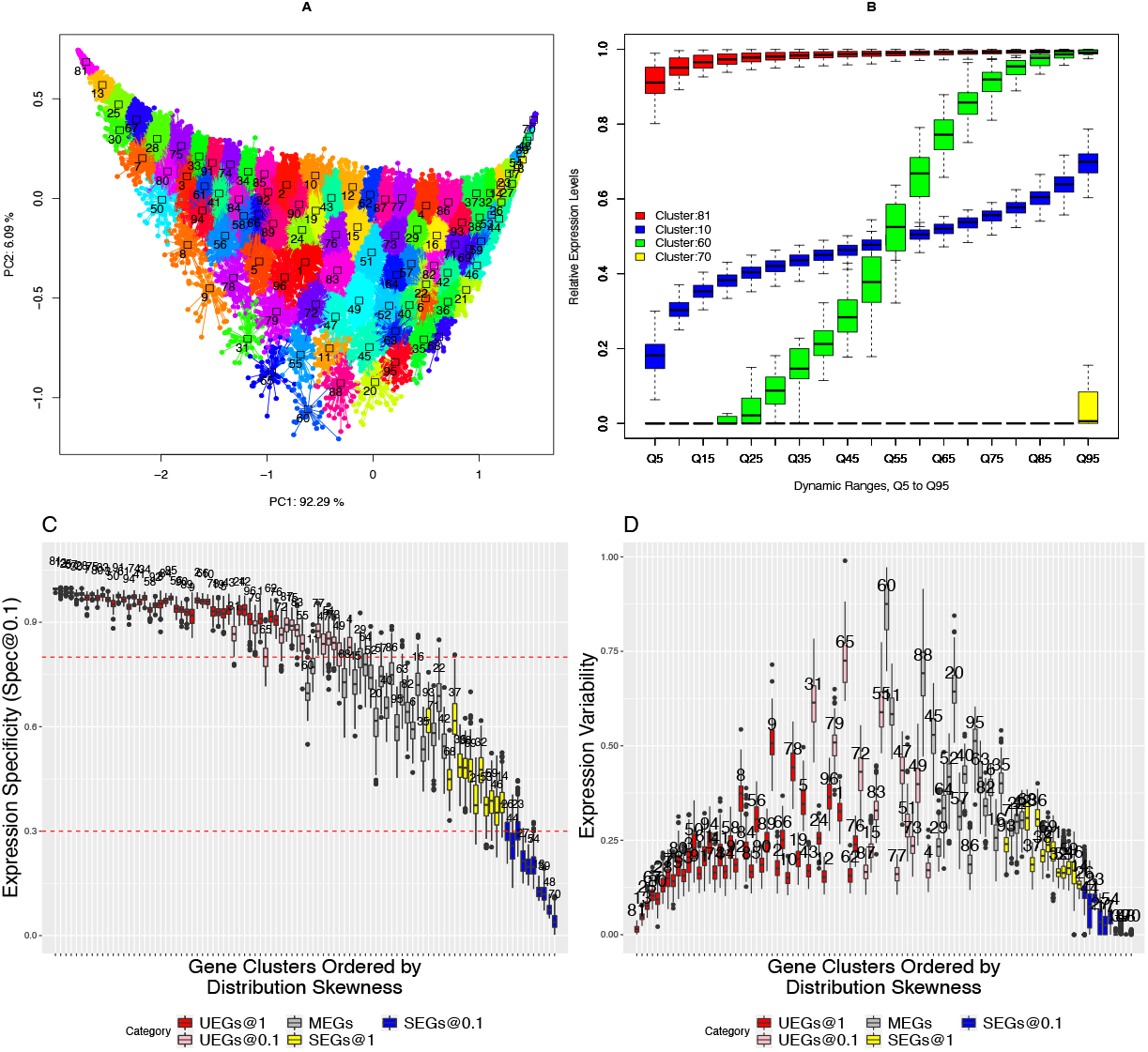
Global Expression Specificity Categories. (A) We use the PCA plot to visualize the global expression patterns and clustering results. we observed that the first PCs (principal components) of the dynamic range matrix correlates the distribution skewness of expression values, and the second PCs correlate the spread of distribution. Each dot represents a gene, and different colors represent 96 gene clusters. (B) The boxplot plot shows the global expression patterns of some gene-clusters. The genes within the same cluster show similar expression levels, expression specificity, and expression variability. (C) The boxplots show the distribution of expression specificity of clusters. The clusters (boxes) are ranked according to their median skewness. The red dashed lines represent the global expression specificity (ϕ) 0.8 (upper) and 0.3 (lower), respectively (using detection threshold of 0.1 TPM). (D) The boxplots show the distribution of expression variability among these clusters. The clusters (boxes) are ranked according to their median skewness.

3) MSG (Moderately Specific Genes) category. This category included 3,150 (12.62%) genes in 20 gene clusters. The genes in this category are mainly involved in regulation of biological processes, including signal transduction (7.59%), cell adhesion (4.66%), and inflammatory response (3.89%).

4) SEG category (SEGs@1) detected by the threshold of TPM 1.0 (with median ϕ of gene clusters <= 0.3). This category included 3,058 (12.25%) genes in 12 clusters.

5) SEG category (SEGs@0.1) only detected by the threshold of TPM 0.1 (with median ϕ of clusters <= 0.3). This category included 6,521 (26.12%) genes in 10 clusters.

The SEG category SEGs@1 and <u>SEGs@0.1</u> refer to those genes that are specifically expressed in a limited set of biological conditions and have specialized functions. The genes in these two specific categories are likely involved in various specific biological processes, such as G-protein coupled receptor signaling pathway (6.58%), sensory perception of smell (4.13%), multicellular organism development (1.88%), and proteolysis (1.72%). All functional enrichment results are listed in Supp Table 9.

### Evaluation the Global Expression Patterns of Disallowed Genes

An interesting example of UEGs associated with vital physiological phenotypes is an important metabolic enzyme gene *Ldha* and a transporter MCT-1 (*SLC16A1*), which belong to a class of so-called disallowed genes, which were first described in the beta cells of pancreatic islets. In contrast to SEGs, disallowed genes refer to those UEGs that are specifically repressed only in a few cell types and with likely functional consequences [26, 27]. For example, the inactivation of *LDHA* and *MCT-1* plays a critical role in the maturation of beta cells and the secretion of insulin. The aberrant activation of *LDHA* or *MCT-1* has been observed to cause diabetes-like phenotype or EIHI (exercise-induced hyperinsulinism). Following the success of *LDHA* and *MCT-1*, a number of putative disallowed genes have been reported [27-29]. Although the repression stability of some putative disallowed genes has been extensively validated [29], they have not been validated from the perspective of UEGs, i.e. the uniqueness of the repression. We think this is partly due to the lack of reliable UEG list and corresponding global expression patterns. The identification and validation of disallowed genes can be viewed as a special application of outlier analysis [30, 31]. Our UEG list provides a resource to evaluate the uniqueness of repression for putative disallowed genes. As shown in Fig 6, the classic disallowed genes *Ldha* and *MCT-1* exhibited strong constitutive expressions across a large collection of transcriptomes. Even for the gene *HK1*, which is specifically repressed in beta cells and liver cells and does not fulfill the strictest definition of disallowance[27], also shows a strong constitutive expression pattern. But some putative disallowed genes exhibited a significantly restricted expression pattern, such as *ITIH5, Cxcl12*, and *HSD11B1*. For example, the *HSD11B1* gene showed a relatively restricted expression pattern in adrenal gland (expressed in 22.58% samples) and bone marrow (expressed 33.70% samples). Besides, although the genes *Igfbp4, MAF, Pdgfra*, and *ARHGDIB* had a ubiquitous expression pattern, their relative expression levels showed significant differences across biological conditions. In addition, these observations were replicated in the Dee2 dataset (Supp Fig 19). These results suggest that unlike the classic disallowed genes, the repression of these genes may be not unique to the islets beta cells.

**Fig 6.**
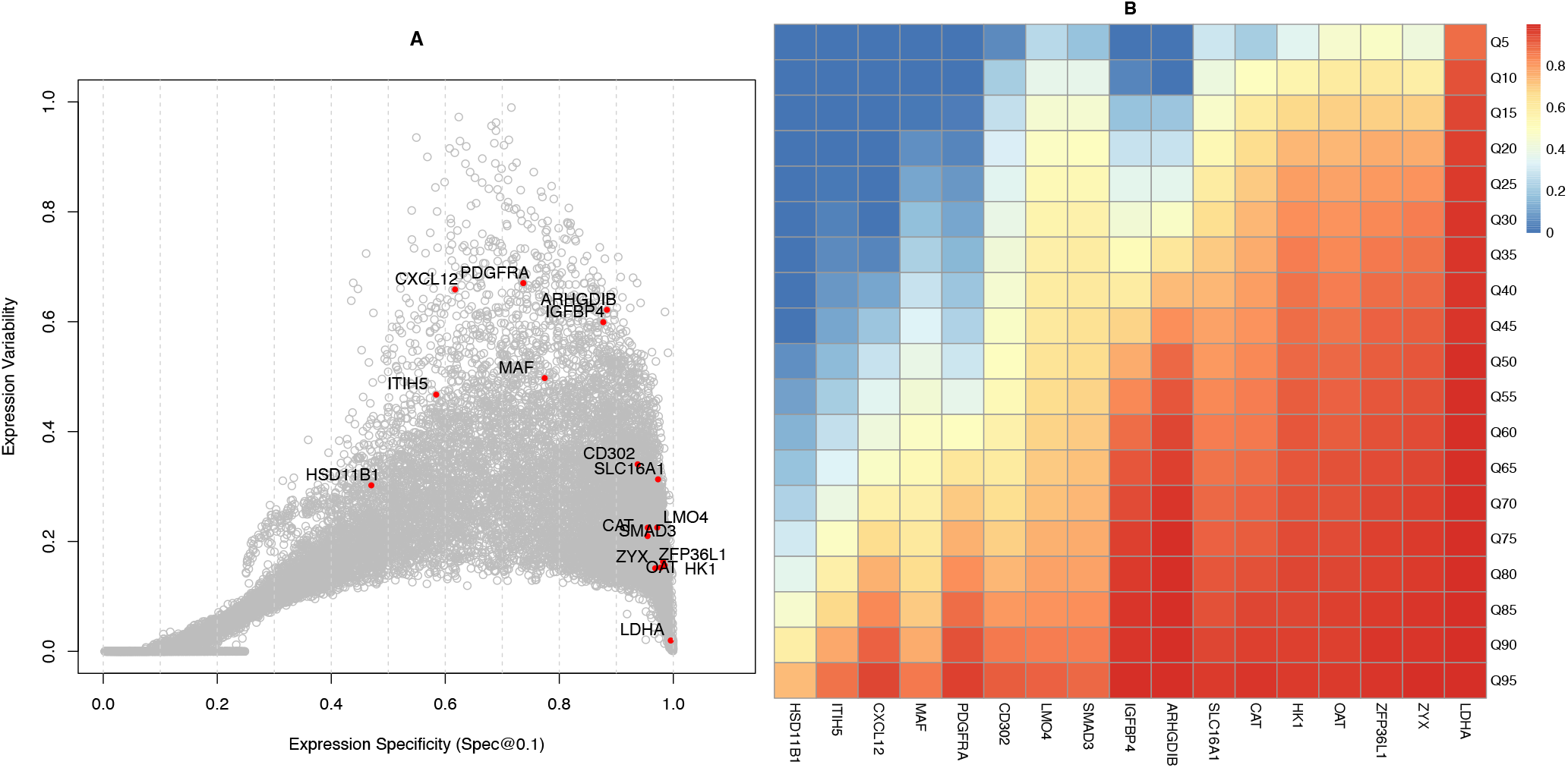
Evaluating the uniqueness of repression for putative disallowed genes in islets beta cells. The classic disallowed gene *Ldha* and *MCT-1* (*SLC16A1*) exhibited a strong ubiquitous expression pattern. The putative disallowed genes *ITIH5*, Cxcl1,2 and *HSD11B1* gene showed a restricted expression pattern, and the *Igfbp4, MAF, Pdgfra*, and *ARHGDIB* genes show relatively high expression variability across biological conditions. These observations suggest that the repression of these seven genes may not be unique to beta cells.

### A large Fraction of UEGs Involves Human Diseases

Since UEGs play an essential role in maintaining cellular processes and cell survival, they were considered unlikely to be a disease gene, especially for genetic diseases [32]. We observed that approximately 80% of the essential genes exhibited a ubiquitous expression pattern (ϕ >= 0.8, Table 1). However, we compared ϕ with the genes associated with physiological traits [33] and genetic diseases [34], and observed that about 70% of physiological traits or diseases related genes exhibited a ubiquitous expression pattern (ϕ >= 0.8, Table 1). For example, loss-of-function mutations in the *ACTB* gene, a most abundant cytoskeletal housekeeping gene, cause development disorder and intellectual disability [35], as well as expanded trinucleotide repeats in the *TBP* gene, an important general transcription initiation factor, cause a Huntington disease-like phenotype [36-38]. Our results indicate that even the most ubiquitously expressed genes cannot be simply ignored during the prioritization of causal genes/variants. On the other hand, genes with restricted expression pattern are believed to be good drug targets due to improved efficacy and safety [4]. This is supported by our observation that about 59.16% of the reported druggable genes [39] show significantly varied ((ϕ < 0.8) expression levels across biological conditions (Table 1).

## Discussion

The goal of our work is to identify and characterize the core genes in the human transcriptome, a long-standing problem in functional genomics. Earlier studies of UEGs relied heavily on the tissue stratification strategy and were limited in sample size, resulting in low consistency across studies and failed to capture global expression patterns, thus limited the potential use of UEG information. As by definition, UEGs should be present across a majority, if not all, phenotypic and physiological conditions of an organism, the laborious and error-prone annotation/curation process may be overcome by the use of a diverse and extensive collection of transcriptomes. In this study, we proposed a global expression specificity metric that uses the proportion of a gene expressed across a large collection of diverse transcriptomes to represent its global expression specificity. Comparisons with results based on tissue specific expression patterns showed that the global expression specificity is highly concordant with tissue specific results (Supp Fig 3-4) and is also robust to uneven distribution of samples across tissue types in the repositories (Supp Fig 6-8).

Leveraging diverse transcriptome profiles, we can establish the global distribution of relative expression values (Fig 3A) for each human gene, and this information can be used to further validate and characterize human UEGs. We examined the relationships between global expression specificity and global distribution attributes of relative expression values. We observed that the UEGs with higher expression levels usually have relatively lower variability in percentile rank within the transcriptome. However, a number of studies found that even for those most commonly used internal reference genes, there was often considerable expression variability across biological conditions [40-43]. About 55.98% of UEGs, especially those highly expressed UEGs, exhibited a narrow distribution of relative expression values, which are correlated with low expression variability (0-0.2) (Supp Table 3). This suggests that most UEGs maintained relatively constant percentile rank within transcriptomes across divergent biological conditions and can serve as good candidates for internal references in most cases. On the other hand, lowly expressed UEGs exhibited relatively higher variability, partly because the percentile ranks of lowly expressed genes are more likely affected by other genes and the size of transcriptomes. In fact, the variability of the observed expression values of a gene is positively correlated with its expression magnitudes (Supp Fig 27).

To better characterize the overall expression patterns for all human genes, we clustered them into clusters, where genes in the same cluster have similar expression levels, expression variability, and expression specificity (Fig 5). With the help of these gene clusters, we identified 19 UEG clusters (LoVarUEGs) containing 5,671 genes with low variability in expression levels. We then checked their dynamic ranges of raw TPM values in both the Recount2 and Dee2 datasets and confirmed their ubiquitous and stable expression patterns (Supp Fig 25 and Supp Table 11-12). After removing outliers (about 3.19%), 5,490 genes had relatively stable TPM values across human transcriptomes and can be used as internal reference genes for expression measurement (Supp Table 13). Compared with previously reported HK genes with stable expression [5], these genes have comparable stability of expression in both the Recount2 and Dee2 datasets. Nevertheless, the LoVarUEGs showed a significantly better coverage for lowly expressed genes (Supp Fig 26). As an advantage over previous studies, our study stratified stably expressed UEGs by their overall expression patterns, so that they can be easily selected and used as internal references for various downstream applications [44]. For example, in this work, we used the lowly expressed UEG clusters to evaluate and determine the optimal expression detection threshold. Interestingly, when we mapped the expression stability of the single-cell [45] onto our gene clusters, we observed that the sparsity (fraction of zeros) of single-cell profiles highly correlates with the global expression specificity and the expression magnitudes at bulk level (Supp Fig 24). The stably expressed UEG clusters with higher expression levels showed lower single-cell sparsity and *vice versa*. This implies that these stably expressed UEG clusters might be a good model to study potential connections in gene expression between bulk and single-cell level, and that maybe useful for cell type deconvolution [46] and adjusting for potential dropouts bias [47]. Moreover, these gene clusters provide local context information for transcriptome profiles that can further improve the outlier analysis approaches [19, 30].

As a validation of our results, we applied our analysis framework to a more recent RNA-seq dataset, Dee2 [17]. As shown in Supp Fig 11, the global expression specificity metric is highly reproducible between these two datasets (Pearson Coefficient: 0.937). Only 5.7% of genes had global expression specificity differences greater than 0.2, and some differences may be caused by different profiling pipelines or gene annotations. In addition, the global distribution attributes for each gene were also highly consistent (Supp Fig 12-13). Finally, a total of 86.2% of UEGs generated from these two repositories overlapped (Supp Fig 15). Comparisons with previous UEG and SEG studies (Table 1) showed that 1) early microarray-based UEG studies significantly underestimated the number of human UEGs; 2) Over 95% of previously reported UEGs were validated in our study (ϕ >= 0.8); 3) we identified 2,804 novel UEGs, 73.57% of which were also found in the separate dataset Dee2 (Supp Fig 15); and 4) there was a significant overlap between UEGs and SEGs. About 37-43% of previously reported SEGs had a strict specific expression pattern (ϕ <= 0.4), but about 21%-28% of these reported SEGs exhibited a ubiquitous expression pattern (ϕ >= 0.8). It implies that some genes may be both ubiquitously and tissue-enriched expressed. For example, the lipid transport gene *APOE*, which is a major risk gene for Alzheimer’s disease [48], showed high expression variability (0.74) while being widely expressed (ϕ = 0.87), and this gene has been labeled as both a UEG or SEG by several studies [3, 8, 9]. The gene *ALAS1*, which is a widely used internal reference gene, was classified as a tissue-specific gene in two recent SEG studies [3, 4]. In addition, we found that even using the same analysis method [3], there was only 38.6% overlap of the identified SEGs between GTEx and BodyMap datasets (Supp Fig 16). Collectively, these observations suggest that a comprehensive study for human SEGs is still required.

Generally, UEGs should be expressed in all living cells of an organism. However, a specific subset of UEGs, called disallowed genes[26], are selectively repressed in some specific cell types with likely functional consequences. The dehydrogenase (*Ldha*) and monocarboxylate transporter-1 (*MCT-1* or *SLC16A1*) genes are the most well-studied disallowed genes in the pancreatic islets beta cells [49]. The repression of *LDHA* is thought to be crucial for the maturation of beta-cells and the secretion of insulin. The beta cells in diabetes models show loss of repression and upregulated expression of *LDHA*. The repression of *MCT-1* prevents the simulation of insulin release inappropriately during physical exercise, and correspondingly, aberrant activating the *MCT-1* gene results in exercise-induced Hyperinsulinism (EIHI) [50, 51]. The identification of disallowed genes in beta cells has raised the interesting question that whether there are other disallowed genes in beta cells or other cell types [49, 52]. Our study provides a comprehensive UEG resource that could be used to evaluate the uniqueness of repression for the identification and validation of disallowed genes. To demonstrate, we evaluated 16 putative disallowed genes in beta cells [27, 29] and found that seven of them (Fig 6 and Supp Fig 19), including *HSD11B1, ITIH5, Cxcl12, Igfbp4, Pdgfra, MAF*, and *ARHGDIB*, exhibited relatively more varied expression patterns. Although our observation is limited to expression level through the UEG perspective, it may offer a new angle for these genes in beta-cells. Moreover, a recent single-cell study revealed that even for the most common UEG genes, such as *GAPDH* and *ACTB*, they showed a clear repression pattern in some cells [45]. This implies that repression of gene expression at single-cell level is likely a common regulatory mechanism and more disallowed genes might exist in specific cell types.

In summary, we have presented a novel data driven framework that uses a large collection of transcriptomes to systematically characterize UEGs. As a major improvement over previous studies, we provide the global expression patterns for human genes that can be used to further validate and characterize UEGs. We also explored some potential functional implications of UEGs in biomedical research and offered an interesting example to demonstrate the usefulness of this resource in the evaluation of disallowed genes.

## Methods

### Pre-processing and Phenotypic Annotation for Transcriptome Profiles

We downloaded 49,649 human transcriptome profiles at the transcript level from the recount2 repository [14, 15]. These profiles were first summarized to gene-level profiles. As the original profiles used Ensembl ID, we converted Ensembl ID into Entrez ID by the Ensembl BioMart Tool (Supp Table 10). If multiple transcripts matched a single Entrez ID, we used the maximum value of these transcripts to represent the expression level of this gene. The gene-level expression matrix was further normalized by TPM (Transcripts Per Kilobase Million) [53] and RPKM (Reads Per Kilobase of transcript, per Million mapped reads). In the following analysis, we found that the TPM threshold had better detection sensitivity so that the main results were analyzed by TPM normalized data and the RPKM-based results were only used for comparison. Because of potential quality issues for transcriptome profiles, we filtered low-quality transcriptome profiles of the Recount2 dataset by the following criterion. If the expression measurement of any of three lowly expressed internal reference genes (*GUSB, HPRT1*, and *HMBS*) in a transcriptome is zero, this transcriptome was considered to be a low-quality profile and excluded (Supp Fig 18). After removal, a total of 39,863 (80.3%) transcriptome profiles remained for further analyses. The Dee2 dataset [17] provided the quality control information for each profile, with a total of 61,020 high-quality Dee2 transcriptome profiles, which were labeled as ‘PASS’, were used for further analysis. We used the same pre-processing method to convert the original profiles into TPM and RPKM normalized matrices.

To check the phenotypic composition of these transcriptomes, each transcriptome was labeled with a series of biomedical ontology terms by an automated semantic annotation database MetaSRA [16], and the sample type was also predicted by MetaSRA. Moreover, we manually annotated 6,501 (16.31%) reference transcriptomes with 101 distinct tissue types and visualized these reference transcriptomes and unclassified transcriptomes by the *onlinePCA* package in R. The manually annotated information of these transcriptomes is listed in Supp Table 6. The overrepresented samples were identified by the PCA ordination density plot with manually determined cutoffs (Supp Fig 2).

### Traditional tissue-based expression specificity and global expression specificity

Traditionally, UEG studies used a tissue stratification strategy to determine the tissue-specificity of expression in order to identify UEGs. This strategy is useful with a limited number of tissue groups and sample size. In this study, we used the proportion of tissues each gene is expressed to represent the traditional tissue-specificity of expression[18]. The manually curated subset was used to calculate tissue specificity for a gene:

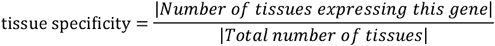

We used TPM 0.1 as the expression detection threshold for each sample. Since there are multiple samples belonging to each tissue group, we used 80% as a cutoff to determine whether this tissue expressed this gene.

However, when one considers a broader spectrum of biological conditions, appropriate grouping samples is non-trivial. It is known that transcriptomes are highly variable across individuals and biological conditions. Therefore, the traditional tissue stratification strategy has hindered the generalization of human UEG study to a larger scale. As UEGs should be broadly expressed in all tissue/cell types of an organism, we propose to use a global expression specificity definition based on the proportion of a gene presented among diverse transcriptomes (Fig 1). This definition does not require defining discrete tissue/cell type groups and is suitable for dealing with a large collection of transcriptomes.

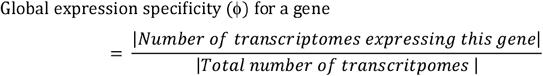

The value of global expression specificity ϕ ranges between 1 for ubiquitously expressed genes and close to 0 for specifically expressed genes.

### Expression Detection Thresholds

With the definition of global expression specificity, the key problem is the appropriate selection of an expression detection threshold to call whether a gene is expressed. We note that there are different methods to define a detection threshold to call a gene expressed [18]. With these lowly expressed UEG clusters, we compared four detection thresholds, with TPM >= 0.1 (which was used in the GTEx Project), TPM >= 1.0, RPKM >= 0.3 [8, 54, 55], and RPKM >=1.0 [56-58]. When using the threshold of TPM 0.1, the median detection rates of low-expressed cluster #4 genes in the Recount2 and Dee2 datasets were 0.83 and 0.89, respectively. However, the detection rates with the threshold of RPKM 0.3 was only 0.65 and 0.58 (Supp Table 13, Supp Fig 25 and Supp Fig 5). We also observed that the detection sensitivity of the RPKM 0.3 was close to that of TPM 1.0. The distribution curves of transcriptome size obtained by RPKM 0.3 and TPM 1.0 showed a significant overlap (Fig 3 B). Altogether, among these commonly used detection thresholds, the detection threshold of TPM 0.1, which was used in the GTEx project, is most sensitive for lowly expressed genes and is more appropriate as the expression detection threshold.

### Gene Functional Annotation and Pathway Enrichment

The manual annotations of the transcriptomes are listed in Supp Table 4. The functional gene sets were downloaded from their original publications and all gene identities were converted to Entrez ID by Ensembl BioMart (Supp Table 10). The gene functional enrichment analyses were conducted by DAVID [52].

#### Quantile Normalization and Batch Effects

To reduce the batch effect and yield a better estimation of global expression patterns, we used a sample-wise quantile transformation to TPM or RPKM normalized expression values for each transcriptome profile [19].

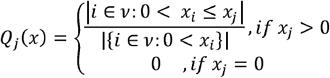

The quantile normalization returns a normalized expression value 0 ≤ *Q*_*j*_ (x) ≤ 1.

After transformation, expression values were replaced by their percentile ranks for each profile. Quantile normalization can eliminate most of the biological and technical variations in expression measurements and result in a semi-quantitative representation for expression levels. We examined the relatedness patterns of quantile normalized profiles by PCA analysis. As shown in Supp Fig 20 and Supp Fig 2, the quantile normalized profiles are highly divergent and reasonably repopulated the entire transcriptome space. We then calculated the within-study differences, within-tissue-group differences, and total differences (Supp Fig 21), and observed that the profiles from the same projects showed relatively higher similarity, but the quantile normalized data significantly reduced the number of outliers. It implies that the quantile normalization method can remove most, but not necessarily all, of the variance attributed to batch. These results suggest that our analysis pipeline may provide a fairly unbiased characterization of gene expression distributions.

### Global Expression Distributions of Relative Expression Values

The sample distribution attributes, including the 5% percentile(Q5) to the 95% percentile (Q95), the interquartile range (IQR), and distribution skewness, of relative expression values, were calculated using R. The series of percentile ranks of relative expression values for each gene (Q5 to Q95), i.e., the dynamic range, represent the global expression pattern for each gene. In this study, we used the IQR (interquartile range) of distribution to represent the expression variability for each gene. Skewness refers to the asymmetry in expression levels. Negative skewness indicates that the most of data points (genes) are towards the high expression end.

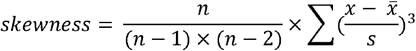

where x is the relative expression value, 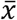 is the sample mean, S is the sample standard deviation, and n is the number of samples.

### Percentile Clustering on Dynamic Ranges of Gene Expression Level

Our goal is to reduce the dimension of the global expression matrix and cluster genes based on their global expression patterns, where members of the same cluster shared similar expression levels, expression variability, and expression specificity (Fig 5). This problem can be formulated as clustering genes by their shapes of distributions. The main difficulty here is the representation of distribution shapes. In this study, we adopted a simple strategy called percentile clustering [23], which uses a series of percentiles of the relative expression values, i.e., the dynamic range matrix, to represent the shape of the distribution, and then uses this percentile matrix to cluster genes. With this strategy, we clustered human genes using an affinity propagation clustering method (apcluster with negDistMat similarity matrices) [59], with the dynamic ranges matrix. Apcluster can infer the number of clusters automatically and provide a representative gene as the local center for each cluster. Our sensitivity analysis reported in supplementary materials shows that the affinity propagation clustering method yielded a better within-cluster homogeneity than the K-means method (Supp Fig 17). Fig 5 A illustrates the clustering results and Supp Table 8 lists the gene clusters.

## Supporting information

SUPPLEMENTARY METERIAL

## Acknowledgements

We thank Dr. Yongkun Wang from the Network and Information Center at Shanghai Jiao Tong University for his support in high-performance computing. We thank PhD Candidate Wei Liu from Yale University for her support in the acquisition of physiological-traits-related genes. HL is supported by the National Key R&D Program of China 2018YFC0910500. JG and JD are supported by the SJTU-Yale Collaborative Research Seed Fund and Neil Shen’s SJTU Medical Research Fund.

## Author Contributions

JG conceived of this study, performed the analysis, and drafted the manuscript. JD contributed to the annotation of metadata and performed the analysis. All authors read and approved the final manuscript. All aspects of the study were supervised by HL and HZ.

