## Supplementary material for "Comprehensive Analysis of Ubiquitously Expressed Genes in Human, From a Data-Driven Perspective": Table 1

### Table 1. The number of genes in each specificity interval.

|  | Global Expression Specificity | | | | | |
| --- | --- | --- | --- | --- | --- | --- |
|  | 0.8-1.0 | 0.6-0.8 | 0.4-0.6 | 0.2-0.4 | 0-0.2 | Total |
| Total Genes | 12,267 (49.14%) | 2,727 (10.92%) | 2,530 (10.13%) | 2,641 (10.58%) | 4,798 (19.22%) | 24,963 |
| Skewness <= 0 | 10,421 (99.2%) | 84 (0.8%) | 0 (0%) | 0 (0%) | 0 (0%) | 10,505 (42.08%) |
| Q10 >= 0.1 | 9,692 (100%) | 0 (0%) | 0 (0%) | 0 (0%) | 0 (0%) | 9,692 (38.83%) |
| Q20 >= 0.1 | 12,002 (99.98%) | 3 (0.02%) | 0 (0%) | 0 (0%) | 0 (0%) | 12,005 (48.09%) |
| 2011 UEGs MICROARRAY | 2,038 (99.66%) | 4 (0.2%) | 2 (0.1%) | 1 (0.05%) | 0 (0%) | 2,045 (8.19%) |
| 2009 UEGs SEQ | 7,703 (98.88%) | 59 (0.76%) | 13 (0.17%) | 13 (0.17%) | 2 (0.03%) | 7,790 (31.21%) |
| 2014 UEGs SEQ | 8,696 (97.6%) | 176 (1.98%) | 37 (0.42%) | 0 (0%) | 1 (0.01%) | 8,910 (35.69%) |
| 2013 HK SEQ | 3,786 (99.82%) | 5 (0.13%) | 2 (0.05%) | 0 (0%) | 0 (0%) | 3,793 (15.19%) |
| BodyMap SEGs | 735 (20.75%) | 573 (16.17%) | 708 (19.98%) | 808 (22.81%) | 719 (20.29%) | 3,543 (14.19%) |
| GTEx SEGs | 1,128 (27.96%) | 661 (16.39%) | 748 (18.54%) | 806 (19.98%) | 691 (17.13%) | 4,034 (16.16%) |
| EssentialGenes | 5,356 (77%) | 548 (7.88%) | 430 (6.18%) | 345 (4.96%) | 277 (3.98%) | 6,956 (27.87%) |
| Traits Genes | 2,246 (72.33%) | 372 (11.98%) | 251 (8.08%) | 172 (5.54%) | 64 (2.06%) | 3,105 (12.44%) |
| GeneticDiseasesGenes | 9,759 (61.42%) | 1862 (11.72%) | 1644 (10.35%) | 1,443 (9.08%) | 1,180 (7.43%) | 15,888 (63.65%) |
| DRUGABLE Genes | 1,764 (40.84%) | 711 (16.46%) | 738 (17.09%) | 643 (14.89%) | 463 (10.72%) | 4,319 (17.30%) |
| UEGs@1 Category | 9,687 (100%) | 0 (0%) | 0 (0%) | 0 (0%) | 0 (0%) | 9,687 (38.81%) |
| UEGs@0.1 Category | 2,272 (89.2%) | 275 (10.8%) | 0 (0%) | 0 (0%) | 0 (0%) | 2,547 (10.20%) |
| MEGs Category | 307 (9.75%) | 2,094 (66.48%) | 749 (23.78%) | 0 (0%) | 0 (0%) | 3,150 (12.62%) |
| SEGs@1 Category | 1 (0.03%) | 358 (11.71%) | 1,771 (57.91%) | 928 (30.35%) | 0 (0%) | 3,058 (12.25%) |
| SEGs@0.1 Category | 0 (0%) | 0 (0%) | 10 (0.15%) | 1,713 (26.27%) | 4,798 (73.58%) | 6,521 (26.12%) |

*SEQ means RNAseq based study. ARRAY means microarray based study.*

*HK is a housekeeping genes study which takes into account the variability of gene expression.*
