## SUPPLEMENTARY METERIAL for "Comprehensive Analysis of Ubiquitously Expressed Genes in Human, From a Data-Driven Perspective"

*Correspondence:

### Supplementary Material List:

**Supplementary Table 1.** Summary of Sample Types.

**Supplementary Table 2.** Summary of Sample Tissue Types.

**Supplementary Table 3.** Number of genes by each variability interval.

**Supplementary Table 4.** Sample types of overrepresentation samples.

**Supplementary Table 5.** Phenotypic composition of overrepresentation samples.

**Supplementary Table 6.** Phenotypic information of transcriptomes.

**Supplementary Table 7.** Global expression specificity and global expression patterns of human genes (Recount2 Quantile Normalized TPM data).

**Supplementary Table 8.** Affinity propagation clustering results.

**Supplementary Table 9.** Functional Enrichment Results.

**Supplementary Table 10.** ID Mapping Table for Human Genes.

**Supplementary Table 11.** The Dynamic Ranges of TPM Values of LoVarUEGs in Recount2 dataset

**Supplementary Table 12.** The Dynamic Ranges of TPM Values of LoVarUEGs in Dee2 dataset

**Supplementary Table 13.** The UEGs Clusters with Stable Expression Magnitude (LoVarUEGs)

**Supplementary Figure 1.** Phenotypic Compositions of Dee2 Samples.

**Supplementary Figure 2.** The overrepresented Level among Analyzed Transcriptomes.

**Supplementary Figure 3.** Global Expression Specificity and Tissue Specificity of Expression.

**Supplementary Figure 4.** 2D Density Plot Between Global Expression Specificity and Tissue Specificity of Expression.

**Supplementary Figure 5.** Comparison of the Global Expression Specificity Categories Identified by Different Normalization Methods and Datasets.

**Supplementary Figure 6.** Estimate the Impacts of Overrepresented Samples on Global Expression Specificity.

**Supplementary Figure 7.** Estimate the Impacts of Overrepresented Samples on Global Distribution Attributes.

**Supplementary Figure 8.** The Differences in Distribution Attributes Between All Samples and Excluded Overrepresented Samples.

**Supplementary Figure 9.** Estimate the Impacts of Sample Size on Global Expression Specificity.

**Supplementary Figure 10.** Estimate the Impacts of Sample Size on Global Distribution Attributes.

**Supplementary Figure 11.** The Reproducibility of Global Expression Specificity Between Two Independent Datasets.

**Supplementary Figure 12.** The Reproducibility of Global Distribution Attributes of Relative Expression Values Between Two Independent Datasets.

**Supplementary Figure 13.** The Differences in Distribution Attributes Between Recount2 and Dee2 Samples.

**Supplementary Figure 14.** Evaluation of the Number of Clusters by Gap-statistics.

**Supplementary Figure 15.** Comparison of UEGs with Previous Studies.

**Supplementary Figure 16.** The SEGs Identified by the Identical Method but Different Datasets Exhibited Significant Discrepancy.

**Supplementary Figure 17.** Comparison of Within-cluster Homogeneity Between Affinity Propagation and K-means Method.

**Supplementary Figure 18.** Transcriptome Profile Quality Control for Recount2 dataset.

**Supplementary Figure 19.** Evaluating the Uniqueness of Repression for Putative Disallowed Genes of the Islets Beta Cells in Dee2 dataset.

**Supplementary Figure 20.** Evaluating the Batch Effects of Transcriptome Profiles, onlinePCA Comparison.

**Supplementary Figure 21.** Evaluating the Batch Effects of Transcriptome Profiles, Compare within-group Similarity.

**Supplementary Figure 22.** The Effect of Gene GC-content on Its Expression Specificity and Expression Pattern.

**Supplementary Figure 23.** 2D-density Plot to Threshold Determination for Quantile Normalized Relative Expression Values.

**Supplementary Figure 24.** Comparison between Single-Cell Expression Stability and Bulk Expression Specificity.

**Supplementary Figure 25.** The Dynamic Ranges of TPM Values of LoVarUEGs by Gene-Clusters.

**Supplementary Figure 26.** Comparison Between LoVarUEGs and 2013HK.

**Supplementary Figure 27.** Comparison of Standard Deviations of TPM Values between LoVarUEGs and 2013HK.

**Supplementary Figure 28.** Comparison of COV of TPM Values between LoVarUEGs and 2013HK.

### Supplementary Results

#### Sensitivity Analysis for Global Expression Specificity and Global Distribution Attributes

As shown in Supp Fig 2, we determined overrepresented sample by 2-D density plot of the first two principal components of onlinePCA. We first checked the phenotypic compositions of these overrepresented samples and observed that they are high divergent in phenotypic compositions (Supp Table 4 - 5). To test if these overrepresented samples would affect global expression specificity and the distribution attributes significantly, we excluded those overrepresented samples and compared their global expression specificity and global distribution attributes with that of all samples. Supp Fig 6 shows that ~90.72% of the genes had difference of global expression specificity less than 0.1 (10% of total range), and the maximal difference was 0.23. For the global distribution attributes (Supp Fig 7-8), the overrepresented samples had a slightly larger impact on the lower bound (Q5) of the distribution of relative expression values, and 16.90% of the genes had their difference larger than 0.1 in Q5. For median relative expression level (Q50), maximal relative expression level (Q95), and expression variability (IQR), more than 90% of the genes had their difference less than 0.1. Collectively, these overrepresented samples had limited effects on the final global expression specificity and distribution attributes. To retain as much information, we used all informative transcriptomes from the recount2 and Dee2 datasets.

Moreover, to evaluate the effects of sample size on the results, we conducted 100 random sampling with a series of sampling sizes for each gene. As shown in Supp Fig 9 and Supp Fig 10, the global expression specificity and distribution attributes became stable as the sample size approached 35,000. These results indicate that we can estimate global expression specificity and global distribution attributes of each gene with a high degree of confidence with a large number of the transcriptomes.

#### Sensitivity Analysis for Percentile Clustering

To validate the clustering robustness and to provide more stable clustering results, we conducted a sensitivity analysis for our percentile clustering results. We assume that the putative variations in clustering results could be largely attributed to 1) the variations in the dynamic range matrix, i.e., the global distribution attributes; 2) the number of clusters; and/or 3) the clustering method.

For the first question, we have discussed in the sensitivity analysis. The results showed that the dynamic range matrix, i.e., global distribution attributes is highly stable and robust. For the second question, we observed that the dynamic ranges of gene expression are continuous (Fig 3A), and it means that there does not seem to be a clear separation boundary in terms of global expression patterns among human genes. We then used the gap-statistic to show the clustering tendency of global expression patterns. The gap statistic compares the sum of within-cluster variations for different numbers of K with their expected values under the null reference distribution of the data. Generally, cluster number with maximum Gap statistic value, which is an elbow point, corresponds to the optimal number of clusters. As shown in Supp Fig 14, as K increased, the Gap statistic showed continuous increase that means the expression patterns do not have a clear optimal number of clusters. To determine the optimal number of clusters or cluster boundaries, we used the affinity propagation clustering method, which can automatically select the optimal number of clusters and does not require the number of clusters to be specified in advance. By comparing clustering results on different datasets and normalization methods, the number of clusters is around 90-96. In this study, we used the clustering results from the recount2 dataset (Supp Table 8, the number of clusters is 96). For the third question, the affinity propagation clustering method is a non-hierarchical clustering method, which simultaneously considers all data points as potential local centers (exemplar), and it requires that all data points within a cluster be similar to its local center (exemplar)[1, 2]. As shown in Supp Fig 17, we made a comparison between affinity propagation and the K-means method using the same cluster number and observed that the affinity propagation method yielded better within-cluster homogeneity (Euclidean distance) than the K-means method.

Collectively, clustering genes by their global expression patterns can better group genes into local homogeneous groups that have similar expression patterns, e.g., expression level, expression variability, and expression specificity. To determine the robustness of clustering results, we validated the expression patterns and global expression specificity of the three clusters in an independent dataset, Dee2. We observed that the gene-clusters identified in the Recount2 data showed similar expression patterns in the Dee2 dataset (Supp Fig 25). We further compared the UEGs categories identified by gene-clusters and observed that the UEGs category is highly reproducible between the Recount2 and Dee2 datasets (86.2% overlapping, Supp Fig 15). These results suggest the effectiveness and robustness of the percentile clustering strategy in this study.

### Supplementary Tables

#### Supp Table 1. Summary of Sample Types.

| Sample Type* | Sample Size |
| --- | --- |
| Tissue | 16,872 (42.32%) |
| Cell line | 13,949 (34.99%) |
| Primary cells | 3,532 (8.86%) |
| In vitro differentiated cells | 3,045 (7.64%) |
| Stem cells | 1,974 (4.95%) |
| Induced pluripotent stem cells | 233 (0.58%) |
| Unknown | 93 (0.23%) |

* Sample type was predicted by MetaSRA database.

#### Supp Table 2. Summary of Sample Tissue Types

| UBERON Term* | Sample Size |
| --- | --- |
| Others | 26,202 (65.73%) |
| Musculoskeletal system | 3,945 (9.90%) |
| Hemolymphoid system | 3,465 (8.69%) |
| Nervous system | 2,978 (7.47%) |
| Digestive system | 1,070 (2.68%) |
| Reproductive system | 666 (1.67%) |
| Immune system | 338 (0.84%) |
| Sensory system | 175 (0.44%) |
| Renal system | 161 (0.40%) |
| Endocrine system | 115 (0.29%) |

* Sematic terms were annotated by MetaSRA database.

#### Supp Table 3. Number of genes by each variability interval.

|  | Global Expression Variability (IQR) | | | | | |
| --- | --- | --- | --- | --- | --- | --- |
|  | 0-0.2 | 0.2-0.4 | 0.4-0.6 | 0.6-0.8 | 0.8-1.0 | Total |
| Total Genes | 15,379 (61.61%) | 7,593 (30.42%) | 1,628 (6.52%) | 316 (1.27%) | 47 (0.19%) | 24,963 |
| Skewness <= 0 | 6,331 (60.27%) | 3,468 (33.01%) | 590 (5.62%) | 105 (1%) | 11 (0.1%) | 10,505 (42.08%) |
| Q10 >= 0.1 | 6,240 (64.38%) | 3,160 (32.6%) | 284 (2.93%) | 8 (0.08%) | 0 (0%) | 9,692 (38.83%) |
| Q20 >= 0.1 | 6,835 (56.93%) | 4,314 (35.94%) | 764 (6.36%) | 91 (0.76%) | 1 (0.01%) | 12,005 (48.09%) |
| 2011 UEGs Microarray | 1,773 (86.7%) | 257 (12.57%) | 11 (0.54%) | 3 (0.15%) | 1 (0.05%) | 2,045 (8.19%) |
| 2009 UEGs SEQ | 5,175 (66.43%) | 2,325 (29.85%) | 254 (3.26%) | 32 (0.41%) | 4 (0.05%) | 7,790 (31.21%) |
| 2014 UEGs SEQ | 5,520 (61.95%) | 2,826 (31.72%) | 426 (4.78%) | 115 (1.29%) | 23 (0.26%) | 8,910 (35.69%) |
| 2013 HK SEQ | 3,212 (84.68%) | 573 (15.11%) | 7 (0.18%) | 1 (0.03%) | 0 (0%) | 3,793 (15.19%) |
| BodyMap SEGs | 2,160 (60.97%) | 993 (28.03%) | 268 (7.56%) | 95 (2.68%) | 27 (0.76%) | 3,543 (14.19%) |
| GTEx SEGs | 2,364 (58.6%) | 1,172 (29.05%) | 349 (8.65%) | 124 (3.07%) | 25 (0.62%) | 4,034 (16.16%) |
| Essential Genes | 4,299 (61.8%) | 2,065 (29.69%) | 479 (6.89%) | 97 (1.39%) | 16 (0.23%) | 6,956 (27.87%) |
| Traits Genes | 1,593 (51.3%) | 1,188 (38.26%) | 264 (8.5%) | 48 (1.55%) | 12 (0.39%) | 3,105 (12.44%) |
| GeneticDiseaseGenes | 8,556 (53.85%) | 5,579 (35.11%) | 1,413 (8.89%) | 294 (1.85%) | 46 (0.29%) | 15,888 (63.65%) |
| Drugable Genes | 2,007 (46.47%) | 1,639 (37.95%) | 504 (11.67%) | 136 (3.15%) | 33 (0.76%) | 4,319 (17.30%) |
| UEGs@1 Category | 6,187 (63.87%) | 3,248 (33.53%) | 246 (2.54%) | 6 (0.06%) | 0 (0%) | 9,687 (38.81%) |
| UEGs@0.1 Category | 662 (25.99%) | 1,127 (44.25%) | 637 (25.01%) | 113 (4.44%) | 8 (0.31%) | 2,547 (10.20%) |
| MEGs Category | 239 (7.59%) | 1,931 (61.3%) | 744 (23.62%) | 197 (6.25%) | 39 (1.24%) | 3,150 (12.62%) |
| SEGs@1 Category | 1,783 (58.31%) | 1,274 (41.66%) | 1 (0.03%) | 0 (0%) | 0 (0%) | 3,058 (12.25%) |
| SEGs@0.1 Category | 6,508 (99.8%) | 13 (0.2%) | 0 (0%) | 0 (0%) | 0 (0%) | 6,521 (26.12%) |

*SEQ means RNAseq based study. ARRAY means microarray based study.*

*HK is a housekeeping genes study which takes into account the variability of gene expression.*

#### Supp Table 4. Sample types of the overrepresentation samples

| Sample Type | Overrepresentation Samples | Total Samples |
| --- | --- | --- |
| Tissue | 7,293 (41.71%) | 16,872 (40.43%) |
| Cell line | 7,967 (45.56%) | 13,949 (34.99%) |
| Primary cells | 563 (3.22%) | 3,532 (8.86%) |
| In vitro differentiated cells | 962 (5.50%) | 3,045 (7.64%) |
| Stem cells | 579 (3.31%) | 1,974 (4.95%) |
| Induced pluripotent stem cells | 53 (0.30%) | 233 (0.58%) |

* Sematic terms were annotated by MetaSRA database.

#### Supp Table 5. Phenotypic composition of overrepresentation samples

| Tissue Type | Overrepresented Samples | Total Samples |
| --- | --- | --- |
| Others | 12,330 (70.52%) | 26,202 (65.73%) |
| Musculoskeletal system | 1,865 (10.67%) | 3,945 (9.90%) |
| Hemolymphoid system | 490 (2.80%) | 3,465 (8.69%) |
| Nervous system | 1,339 (7.66%) | 2,978 (7.47%) |
| Digestive system | 311 (1.78%) | 1,070 (2.68%) |
| Reproductive system | 453 (2.59%) | 666 (1.67%) |
| Immune system | 79 (0.45%) | 338 (0.84%) |
| Sensory system | 115 (0.66%) | 175 (0.44%) |
| Renal system | 47 (0.27%) | 161 (0.40%) |
| Endocrine system | 42 (0.24%) | 115 (0.29%) |

* Sematic terms were annotated by MetaSRA database.

#### Supp Tables 6-13 are provided as separate files.

### Supplementary Figures.

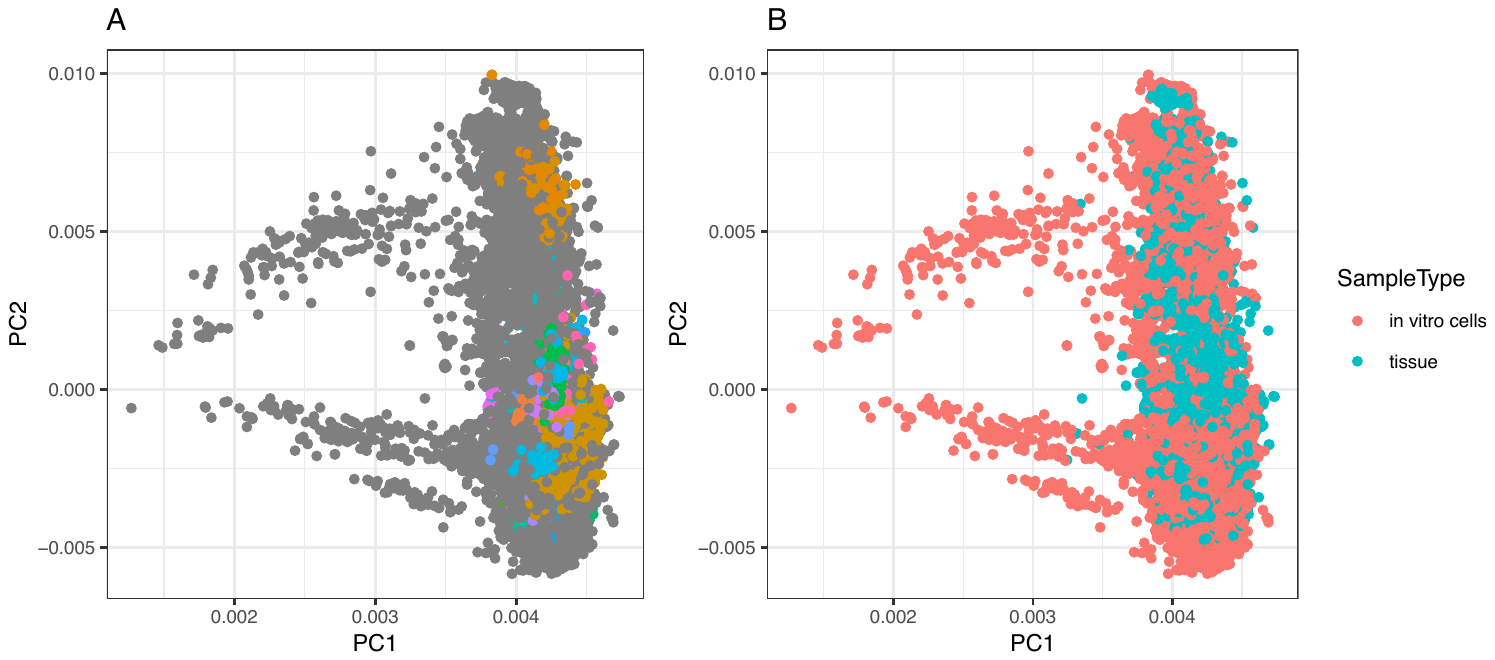

#### Supp Fig 1. Phenotypic Compositions of Dee2 Samples.

The onlinePCA was performed to the quantile normalized TPM data to visualize the phenotypic compositions and relatedness among transcriptomes for the Dee2 dataset. Each dot represents one transcriptome projected on the principal plane formed by the first and second principal axes.

(A). The colored dots represent the 6,501 (16.31%) manually curated reference transcriptomes belonging to 101 tissue groups. Dark dots represent those unclassified transcriptomes that exhibited a wide spectrum of heterogeneity.

(B). The blue dots are the transcriptomes from tissue samples, and the red dots are the transcriptomes from in vitro cells.

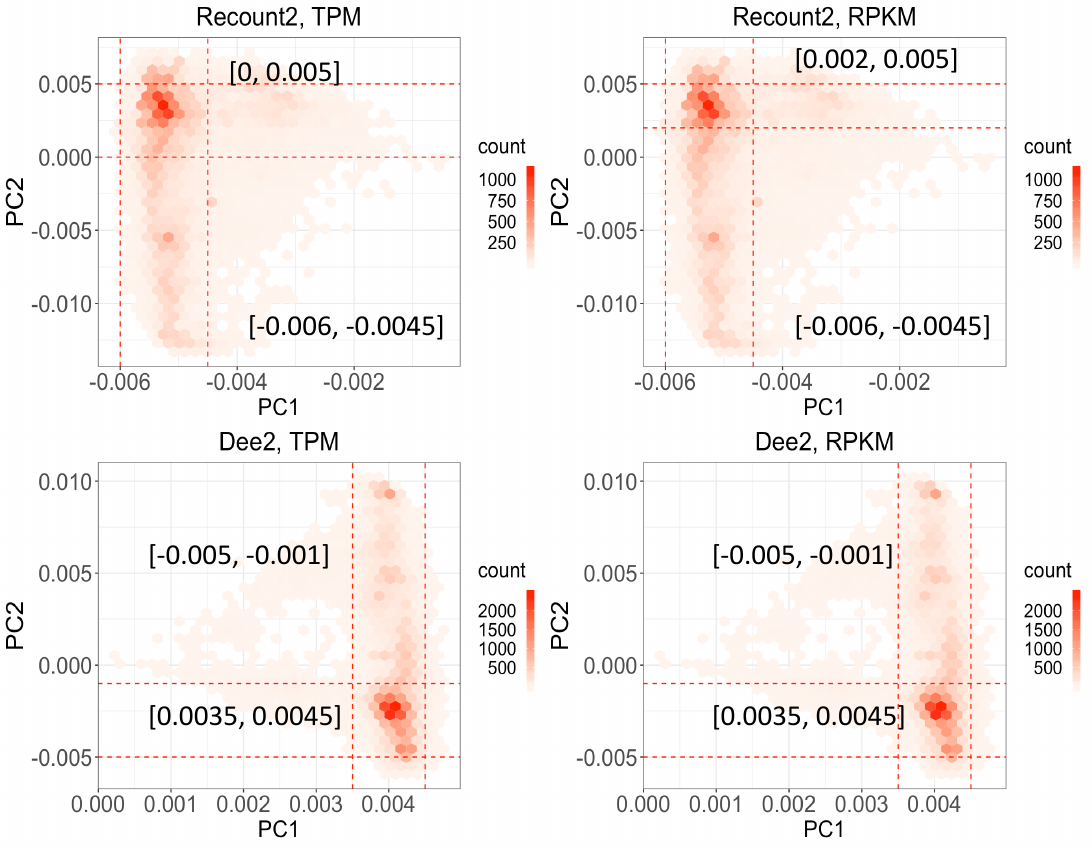

#### Supp Fig 2. The Overrepresented Level among Analyzed Transcriptomes.

PCA ordination density plot of the first and second principal components. The color of each hexagon represents the corresponding sample density. The dashed area indicates these overrepresented samples.

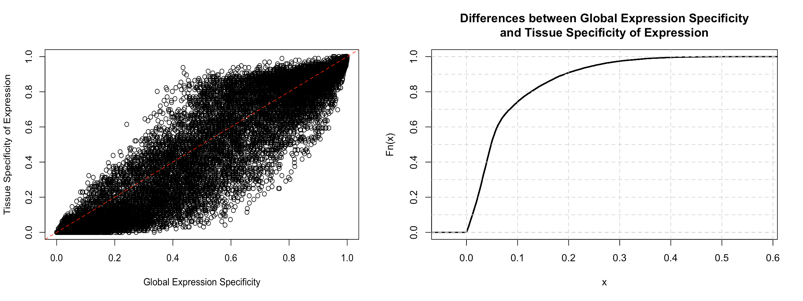

#### Supp Fig 3. Global Expression Specificity and Tissue Specificity of Expression.

The left figure is the comparison between global expression specificity and traditional tissue specificity of expression. The right figure is the cumulative distribution curve of the absolute differences between these two metrics. The expression detection threshold is TPM 0.1. We observed that the global expression specificity highly concordant with traditional tissue specificity of expression (Pearson Coefficients is 0.960) and only 2,279 genes (9.1%) that have relatively high divergence (>=0.2) between these two metrics.

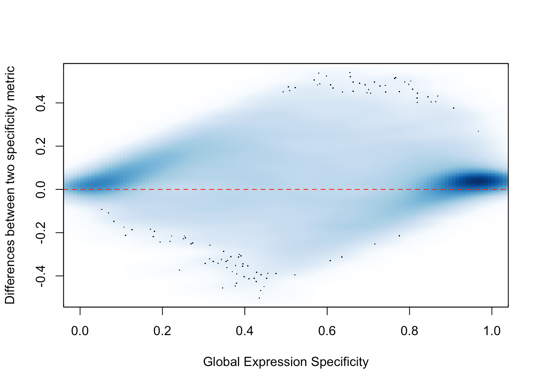

#### Supp Fig 4. 2D Density Plot Between Global Expression Specificity and Tissue Specificity of Expression.

The 2D density plot show that the genes with higher or lower global expression specificity have higher agreement between global expression specificity and traditional tissue specificity of expression.

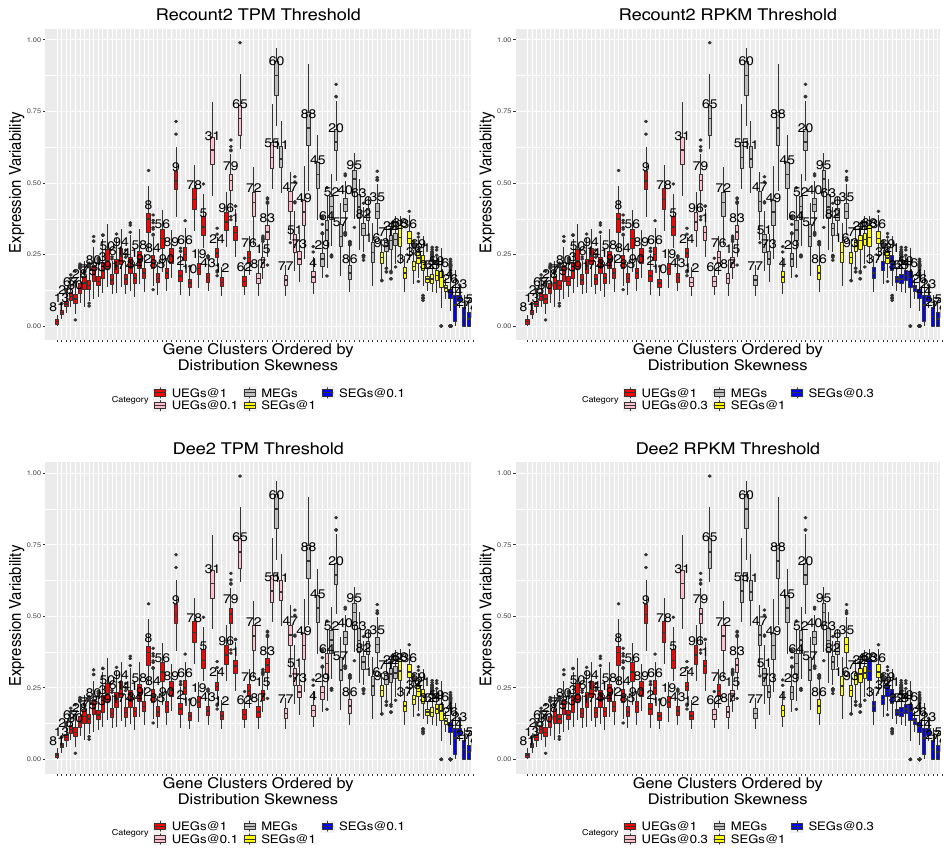

#### Supp Fig 5. Comparison of the Global Expression Specificity Categories Identified by Different Normalization Methods and Datasets.

We compared the global expression categories between different expression detection thresholds, including TPM 0.1, TPM 1.0, RPKM 0.3 and RPKM 1.0. The global expression patterns and gene clusters were obtained from Recount2 quantile normalized TPM data. The detection threshold of TPM 0.1 identified more UEGs genes than RPKM 0.3.

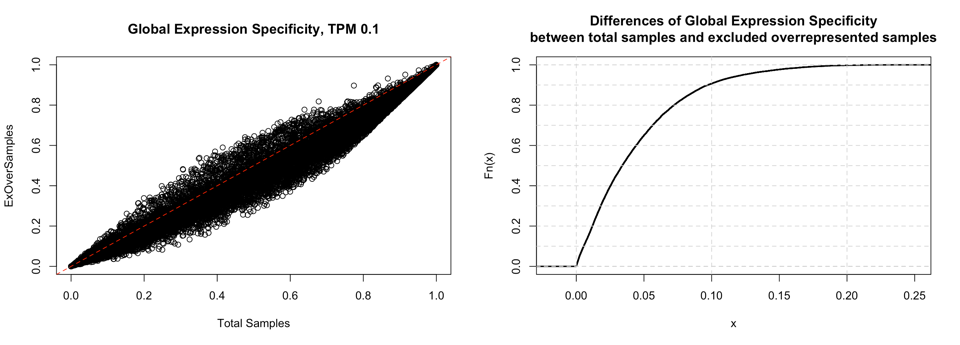

#### Supp Fig 6. Estimate the Impacts of Overrepresented Samples on Global Expression Specificity.

The left figure is the scatter plot of the global expression specificity obtained from total informative samples of the Recount2 dataset and the samples which excluded the overrepresented samples. They are highly concordant with each other (Pearson coefficient is 0.99). The right figure is the cumulative distribution curve of the absolute differences of the global expression specificity between them and about 90.72% of total genes, have the difference of global expression specificity was less than 0.1 (10% of total range), and the maximal difference is 0.23. The Global Expression Specificity determined by the threshold of TPM 0.1.

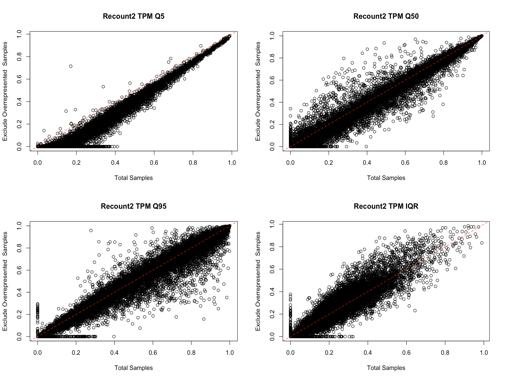

#### Supp Fig 7. Estimate the Impacts of Overrepresented Samples on Global Distribution Attributes.

These figures show the 4 major distribution attributes between total informative samples of the Recount2 dataset and the samples which excluded the overrepresented samples. We observed that the overrepresented samples only have a slightly larger impact on the lower bound (Q5) of the distribution of relative expression values.

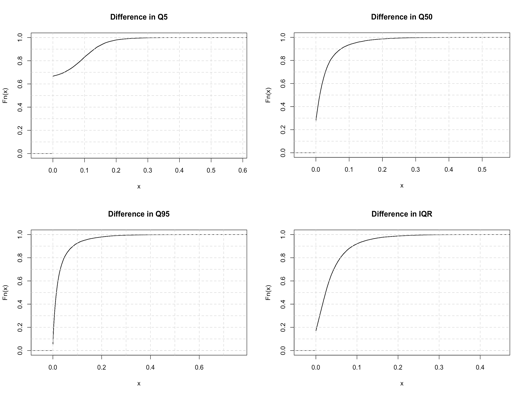

#### Supp Fig 8. The Differences in Distribution Attributes Between All Samples and Excluded Overrepresented Samples.

These figures show the differences of 4 major distribution attributes between total informative samples of the Recount2 dataset and the samples which excluded the overrepresented samples. We observed that about 16.90% of the genes had their difference larger than 0.1 in Q5. For median relative expression level (Q50), maximal relative expression level (Q95), and expression variability (IQR), more than 90% of the genes had their difference less than 0.1.

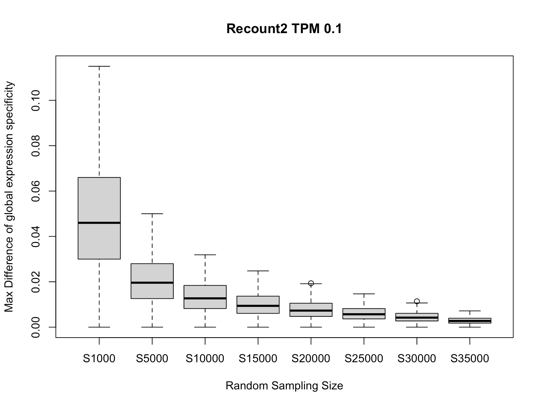

#### Supp Fig 9. Estimate the Impacts of Sample Size on Global Expression Specificity.

We conducted 100 times random sampling for each gene and a series of sampling size (x-axis), including 1000, 5000, 10000, 15000, 20000, 25000, 25000, 30000, and 35000. The global expression specificity become stable as the sampling size approaches 35,000. The median maximal differences of global expression specificity is 0.0027 (0.25% of total range).

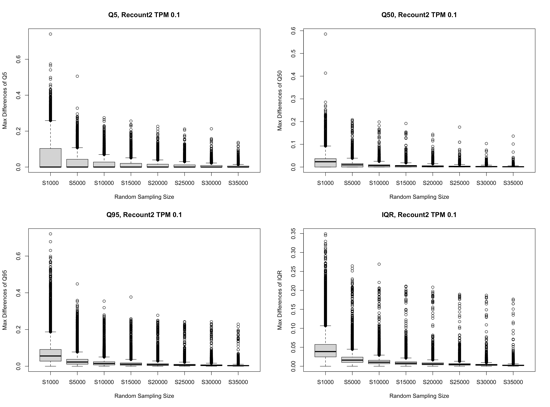

#### Supp Fig 10. Estimate the Impacts of Sample Size on Global Distribution Attributes.

We conducted 100 times random sampling for each gene and sampling size (x-axis), including 1000, 5000, 10000, 15000, 20000, 25000, 25000, 30000, and 35000. The 4 major distribution attributes become stable as the sampling size approaches 35,000. The median maximal differences of these attributes are less than 0.004 (0.4% of total range).

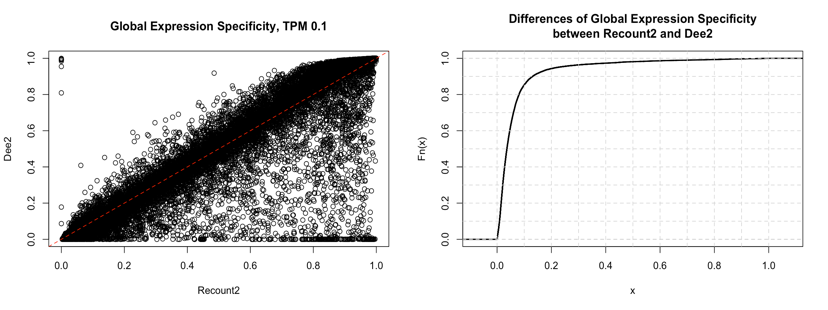

#### Supp Fig 11. The Reproducibility of Global Expression Specificity Between Two Independent Datasets.

The left figure is the scatter plot of the global expression specificity obtained from Recount2 and Dee2, respectively. The right figure is the cumulative distribution curve of the absolute differences in the global expression specificity between these two datasets. The Global Expression Specificity determined by the threshold of TPM 0.1.

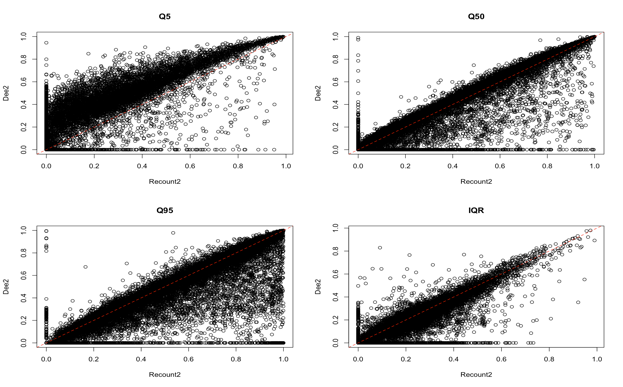

#### Supp Fig 12. The Reproducibility of Global Distribution Attributes of Relative Expression Values Between Two Independent Datasets.

These figures show the 4 major distribution attributes obtained by Recount2 and Dee2 datasets.

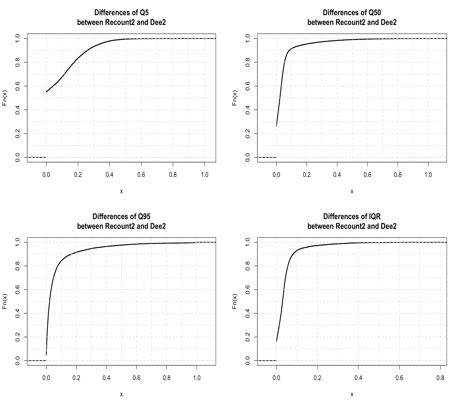

#### Supp Fig 13. The Differences in Distribution Attributes Between Recount2 and Dee2 Samples.

These figures show the differences of 4 major distribution attributes between Recount2 of the Dee2 samples. We observed that about 16.61 % of the genes had their difference larger than 0.2 in Q5. For median relative expression level (Q50), maximal relative expression level (Q95), and expression variability (IQR), more than 90% of the genes had their difference less than 0.2.

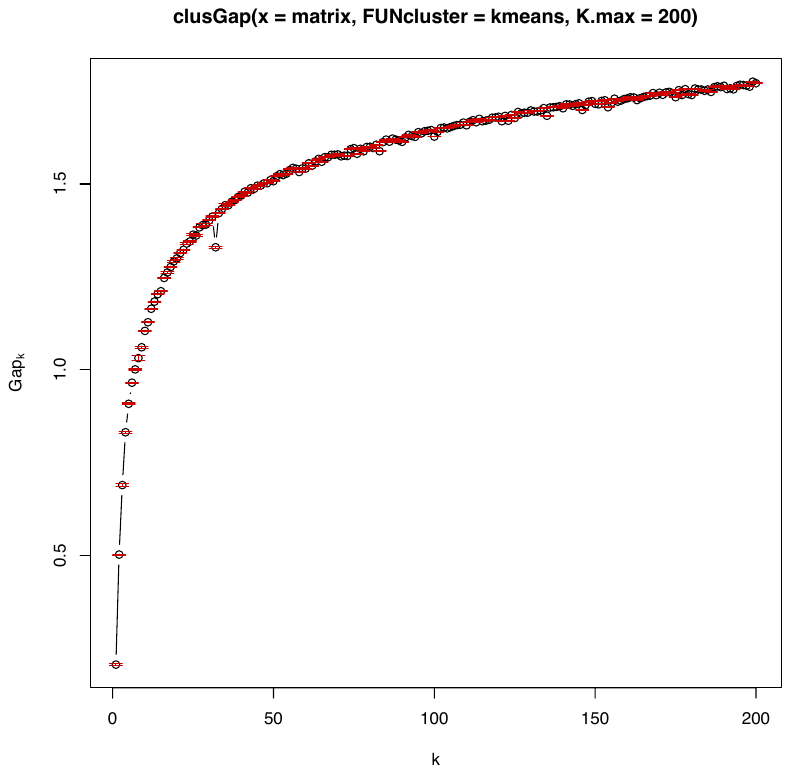

#### Supp Fig 14. Evaluation of the Number of Clusters by Gap-statistics.

Generally, cluster number with maximum Gap statistic value corresponds to optimal number of clusters. However, as K increases, the Gap statistics shows continuous and smooth growth. We did not observe a clear elbow point. The quantile normalized TPM data with K-means clustering method used to calculate this gap-statistics.

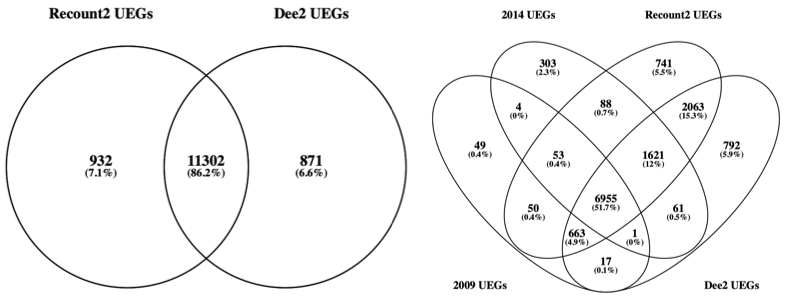

#### Supp Fig 15. Comparison of the UEGs Category with Previous Studies.

Comparisons with previous UEGs and SEGs studies (Table 1) showed that 1) early microarray-based UEG studies significantly underestimated the number of human UEGs; 2) Over 95% of previously reported UEGs were validated in our study (φ >= 0.8); 3) we identified 2,804 novel UEGs, 73.57% of which were also found in the independent dataset Dee2; 4) a total of 86.2% of UEGs generated from Recount2 and Dee2 overlapped.

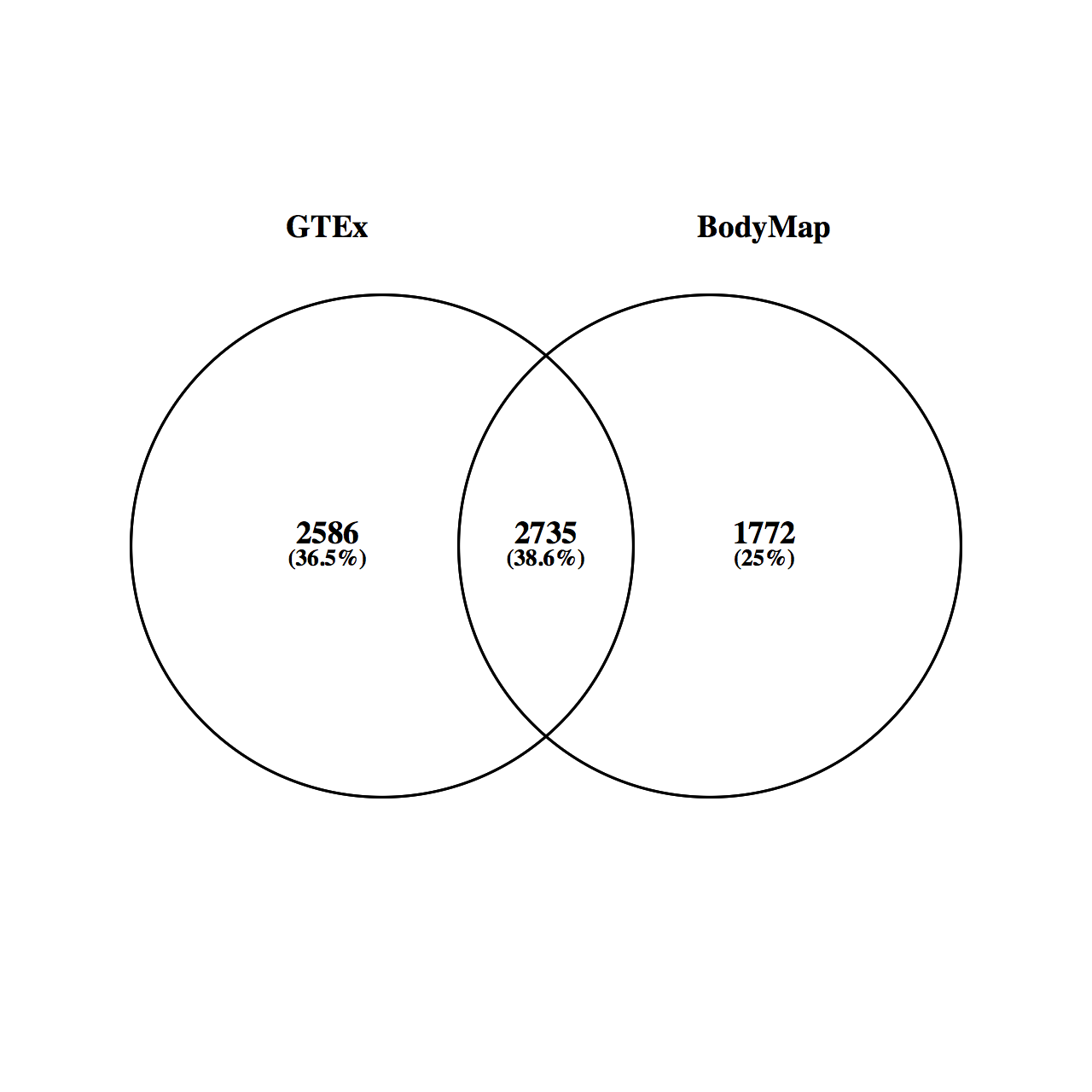

#### Supp Fig 16. The SEGs Identified by the Identical Method but Different Datasets Exhibited Significant Discrepancy.

Even using the same method, the specifically expressed genes identified by different datasets still showed considerable levels of discrepancy. These two gene sets downloaded from a recent SEGs study [PMID: 29309507].

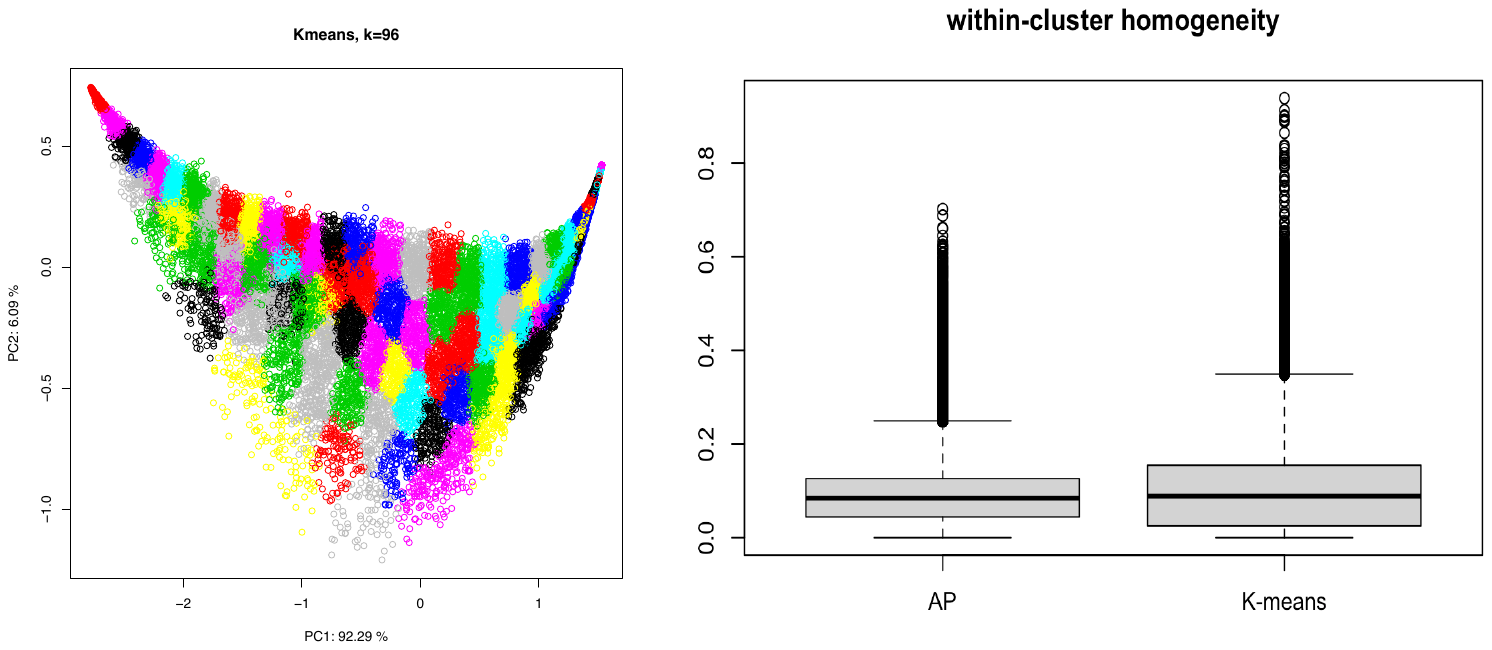

#### Supp Fig 17. Comparison of within-cluster Homogeneity Between Affinity Propagation and K-mean Method

The left figure is the clustering results of K-means method with the sample cluster number (96) of affinity propagation (AP) method. The right figure shows that AP method yielded better within-cluster homogeneity than the K-means method.

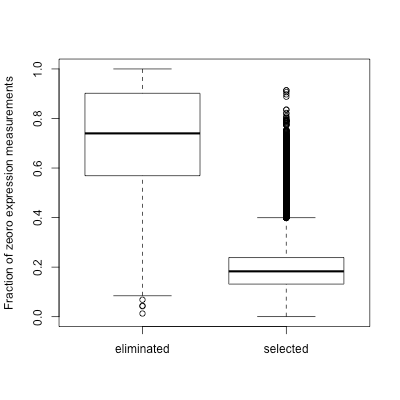

#### Supp Fig 18. Transcriptome Profile Quality Control for Recount2 Dataset.

A transcriptome was considered as a low-quality profile if any of 3 low-expression internal reference genes (*GUSB*, *HPRT1*, and *HMBS*) had expression measurements of zero, and were eliminated for further analysis. We used boxplot to compare the sparsity level between the low-quality profiles and the informative profiles.

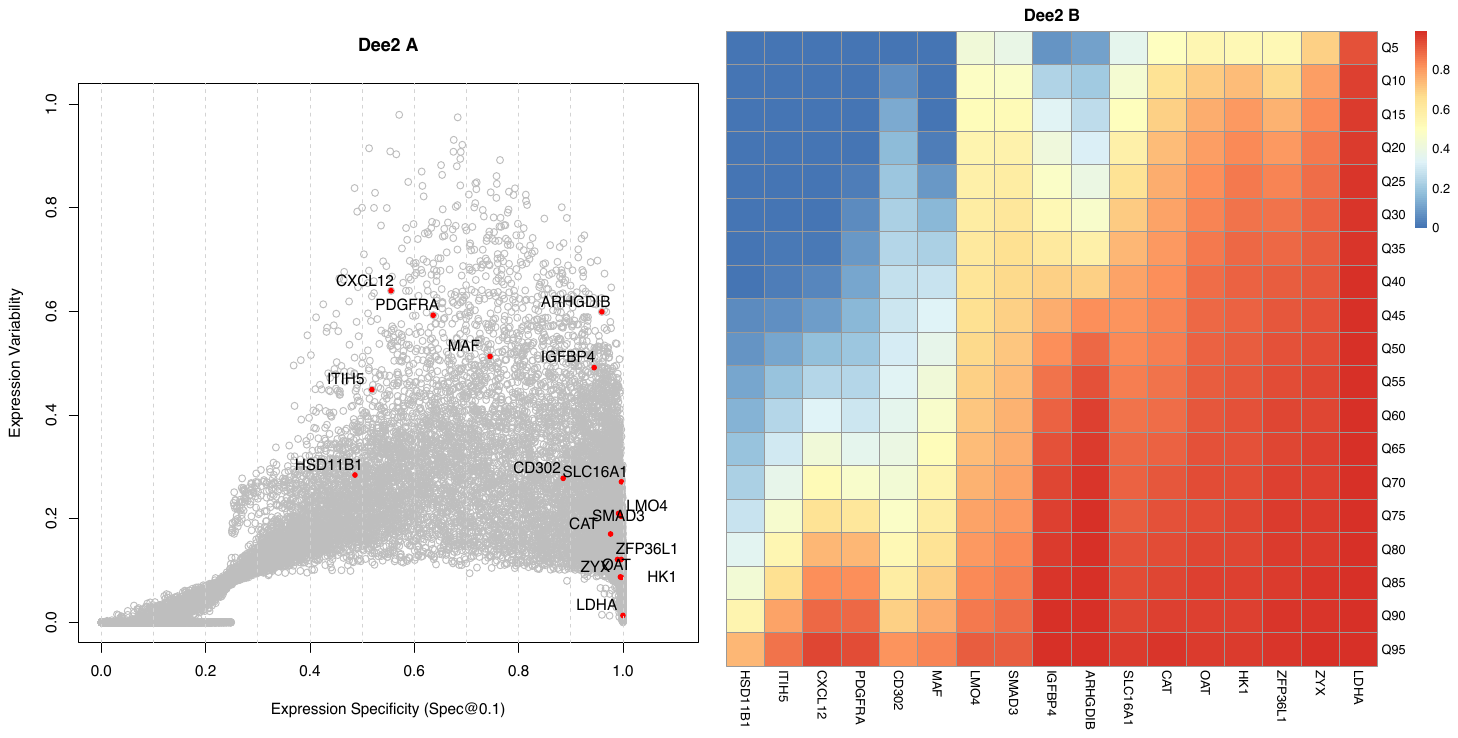

#### Supp Fig 19. Evaluating the Uniqueness of Repression for Putative Disallowed Genes of The Islets Beta Cells in Dee2 dataset.

The global expression specificity and global expression patterns of the 16 putative disallowed genes are consistent with the observations in the Recount2 dataset.

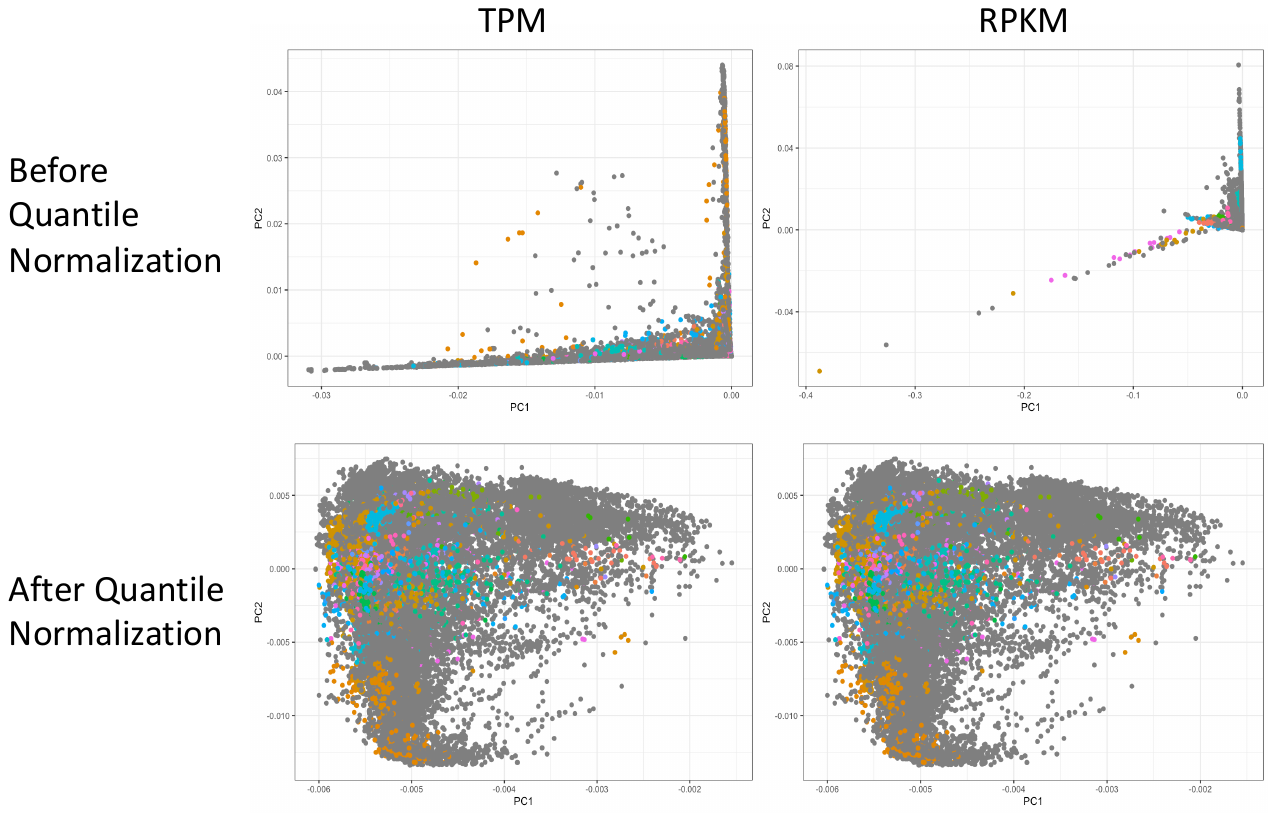

#### Supp Fig 20. Evaluate the Batch Effects of Transcriptome Profiles, onlinePCA Comparison.

After quantile normalization, we observed that the data points (Recount2 samples) reasonably repopulate the entire transcriptome space. It implies the quantile normalized data significantly reduced the batch effects.

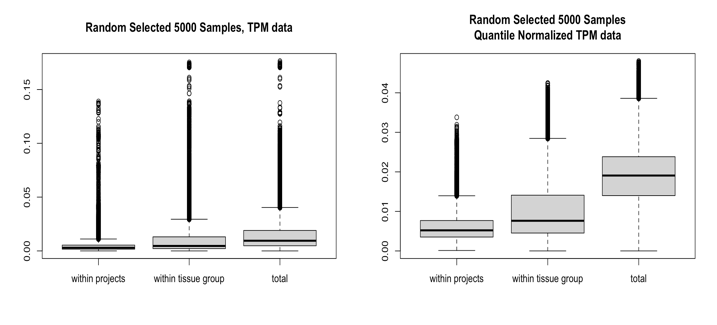

#### Supp Fig 21. Evaluate the Batch Effects of Transcriptome Profiles, Compare within-group Similarity.

We randomly selected 5000 profiles and calculated the Euclidean distance between data points that within-tissue-group, within-projects, and total background. We observed that the profiles from the same projects show relative higher similarity, but the quantile normalized data significantly reduced the number of outliers. It implies that the quantile normalization method can remove most, but not necessarily all, of the variance attributed to batch.

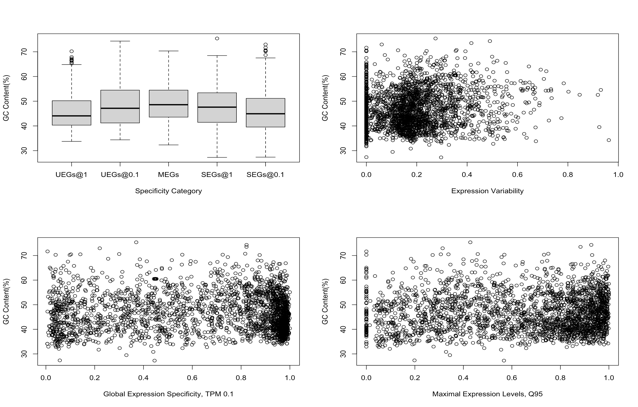

#### Supp Fig 22. The Effect of gene GC-content on Its Expression Specificity and Expression Pattern.

We did not observe any significant relationships between GC content and its expression specificity or expression pattern. The GC-content information was obtained from the Ensembl BioMart.

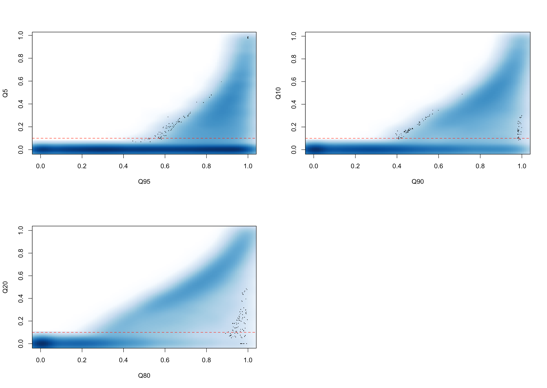

#### Supp Fig 23. 2D-density Plot to Threshold Determination for Quantile normalized Relative Expression Values.

The dashed red line is the relative expression level at 0.1 that could be used to empirically distinguish expression genes from non-expressed genes.

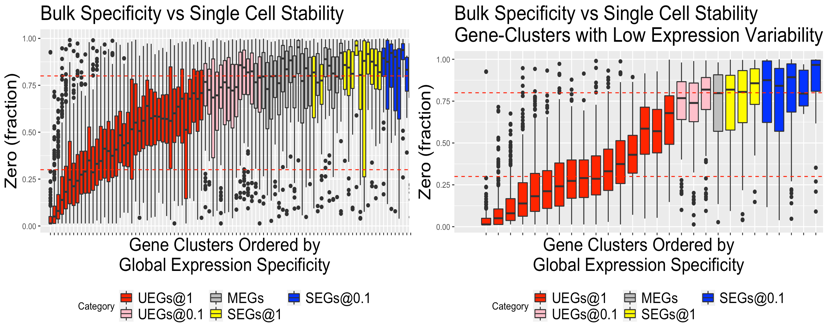

#### Supp Fig 24. Comparison between Single-Cell Expression Stability and Bulk Expression Specificity

We mapped the expression stability of the single-cell onto our gene-clusters and observed that the sparsity (fraction of zeros) of single-cell profiles highly correlates with the global expression specificity and the expression magnitudes at bulk level. Left figure is all 96 gene clusters ranked by their median global expression specificity. Right figure is the 19 UEGs clusters with low expression variability, including Cluster #81, #13, #25, #67, #28, #75, #33, #91, #74, #34, #85, #2, #10, #43, #12, #62, #87, #77 and #4, and 9 MEGs/SEGs clusters with low expression variability #70, #48, #39, #54, #23, #14, #32, #37 and #86. (From left to right in the right figure)

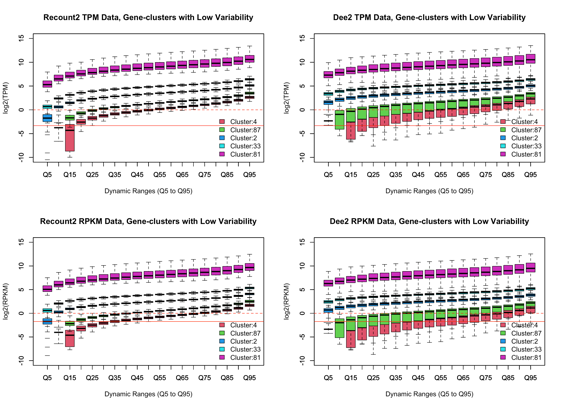

#### Supp Fig 25. The Dynamic Ranges of TPM Values of LoVarUEGs by Gene Clusters

We checked the dynamic ranges of raw TPM values for some LoVarUEGs gene clusters in both of Recount2 and Dee2 dataset, thus confirmed their ubiquitous and stable expression patterns. Dashed lines in the upper two figures are the threshold of TPM 1.0 and solid lines are the threshold of TPM 0.1. Dashed lines in the lower two figures are the threshold of RPKM 1.0 and solid lines are the threshold of RPKM 0.3.

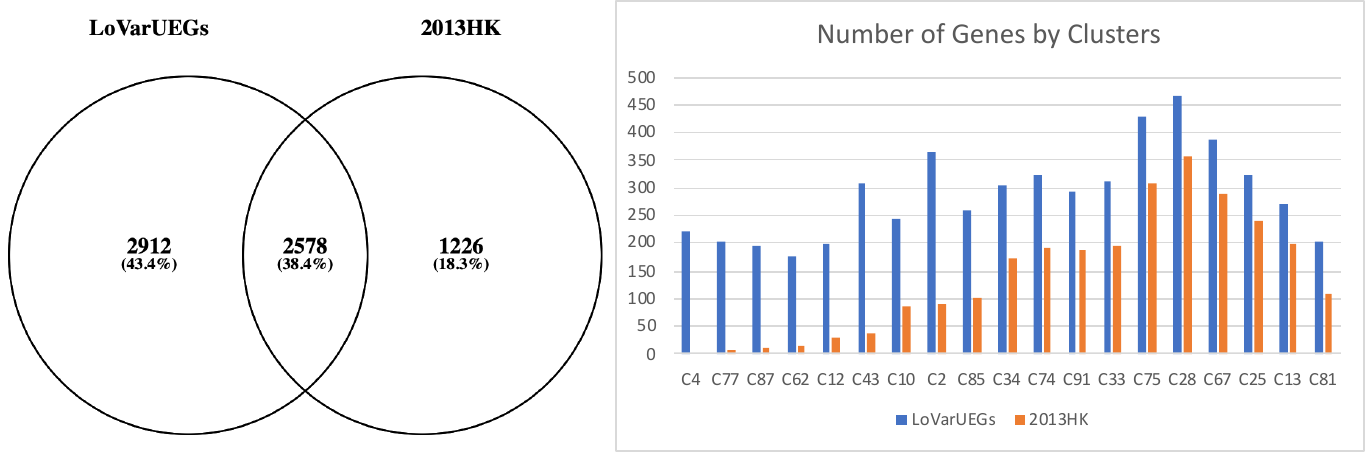

#### Supp Fig 26. Comparison Between LoVarUEGs and 2013HK

Compared with the previous reported HK genes with stable expression, we observed that the LoVarUEGs shows a significantly better coverage for those low expressed genes. x-axis is the gene clusters ordered by expression magnitude. Left figure is the venn diagram between 2013HK and LoVarUEGs. Right figure is the number of genes by gene-clusters. x-axis is the gene-clusters. The LoVarUEGs is provided in Supp Table 13.

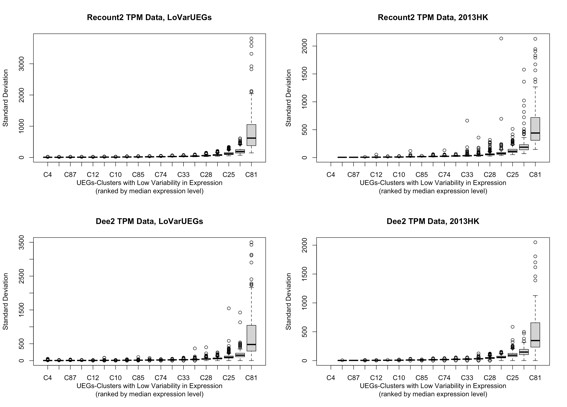

#### Supp Fig 27. Comparison of Standard Deviations of TPM Values between LoVarUEGs and 2013HK

Compared with the previous reported HK genes with stable expression, we observed that they have comparable standard deviations of expression in both of Recount2 and Dee2 dataset. The LoVarUEGs is provided in Supp Table 13 and their dynamic ranges is provided in Supp Table 11-12.

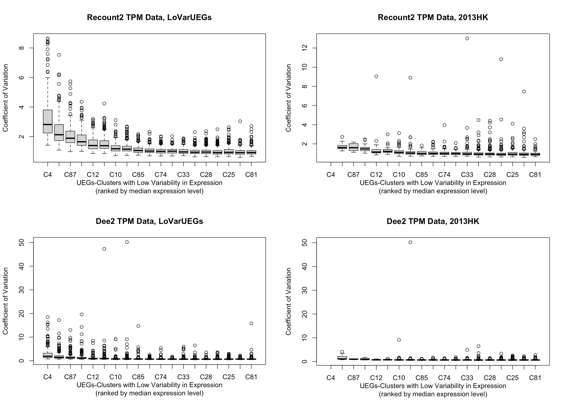

#### Supp Fig 28. Comparison of COV of TPM Values between LoVarUEGs and 2013HK

Compared with the previous reported HK genes with stable expression, we observed that they have comparable COV (coefficient of variations) of expression in both of Recount2 and Dee2 dataset.
